# Dietary serine and glycine restriction delays cancer-associated cachexia

**DOI:** 10.64898/2026.10.15.759471

**Authors:** Vassilis Papalazarou, Jack Coomes, Martha M. Zarou, Izadora Furlani, Karen H. Vousden

## Abstract

Cancer cachexia is a severe metabolic complication of malignancy characterized by loss of appetite, progressive tissue wasting leading to adipose tissue and/or muscle loss and systemic inflammation. Energy dysregulation and nutrient availability are increasingly recognized as key determinants of host metabolic adaptation during tumour progression. However, how diets can be used to support the host remains unclear and conventional dietary regimes have failed to reverse cancer cachexia. Here, we show that dietary serine/glycine restriction induces a coordinated host metabolic reprogramming that delays the onset of cachexia in tumour-bearing mice. Healthy mice fed a serine/glycine free diet show modest body weight loss resulting from reduced caloric intake and reduced adiposity. Surprisingly, in cancer bearing mice, the remodelled metabolic state leads to preservation of adipose depots and delayed cancer associated cachexia. This response is independent of the impact of serine availability on tumour growth, suggesting that it is the result of host-driven adaptations rather than a direct reflection of tumour burden. Mechanistically, we show that serine and glycine restriction remodels systemic metabolism, leading to reduced oxidative lipid stress and attenuation of inflammatory responses. Together, these findings identify serine/glycine availability as a key regulator of host metabolic resilience during cancer progression. These data support a model in which dietary amino acid restriction modulates systemic inflammation and lipid metabolism to delay the onset of cachexia, highlighting potential therapeutic avenues aimed at targeting host metabolic state rather than tumour growth alone.

**Graphical Abstract:** 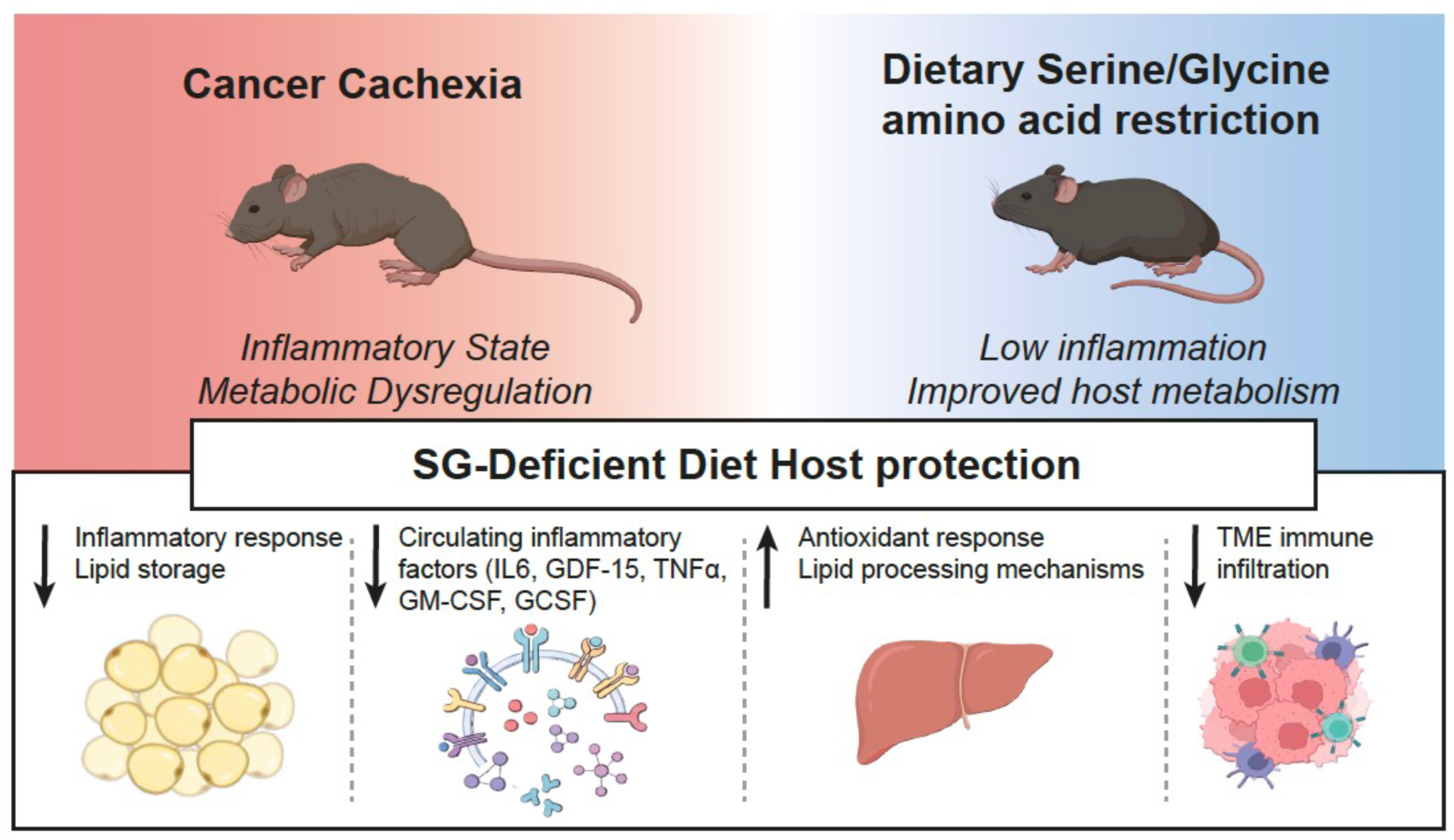

## INTRODUCTION

Dietary interventions have emerged as a promising strategy to exploit metabolic liabilities in cancer and are becoming increasingly refined with our growing understanding of the mechanisms through which cancer cells rewire their metabolism^1–3^. Several metabolic vulnerabilities associated with malignant transformation have been identified^4–6^, including increased dependence of cancer cells on serine metabolism and flux through the one carbon cycle^7,8^. The potential for manipulation of this metabolic pathway is highlighted by the success of anti-folates as one of the earliest successful targeted chemotherapies^9,10^. Numerous studies have shown that dietary depletion of serine and glycine can have a substantial impact on the circulating levels of these two amino acids, despite the ability of most cells to synthesise serine *de novo* from the glycolytic intermediate 3-phosphoglycerate, with dietary serine/glycine limitation being effective in delaying tumour growth and extending survival in many, but not all cancer models ^11–17^.

While dietary interventions are an attractive approach for cancer patients, the potential for adverse reactions to dietary modulation of specific nutrients remains a concern. Normal cells also require serine for growth and rapidly proliferating cells such as activated T cells have been shown to depend on serine or other components of serine metabolism, such as formate, for function^18–20^. However, a serine/glycine deficient diet has also been demonstrated to promote antitumor immunity and appeared a safe therapeutic strategy in a phase I clinical trial^21^. Certain cell types, including hematopoietic stem cells (HSCs) depend on extracellular serine uptake and serine/glycine-deficient diet affected HSC regeneration following irradiation-induced myelosuppressive injury in mice^22^. Long term low systemic serine levels are also associated with certain pathologies, including macular disease and peripheral neuropathy^23^ as well as diabetes-associated neuropathy^24^. Specifically, extended dietary serine/glycine restriction in combination with a high fat diet can lead to small fibre neuropathy and reduced adiposity^24^. Taken together, while serine/glycine free diets appear to be well tolerated in the short term, extended use may lead to undesirable responses. Further research is required to assess the impact of serine depletion on therapeutic response.

Previous work has shown that dietary serine and glycine depletion combined with small molecule inhibition of the *de novo* serine synthesis pathway led to weight loss, although this was readily reversed by modulation of the dose of the inhibitor, and these mice remained healthy without obvious evidence of brain, liver or gut toxicity^14^. The serine/glycine deficient diet alone led to a modest but limited weight loss in mice^13,14,24^.

Many cancers, including pancreas and lung tumours, are associated with an involuntary loss of body weight, associate with fat and/or muscle wasting, in a syndrome known as cancer associated cachexia^25^. Cachexia results in a significant decrease in quality of life, including disrupted sleep, lower activity and anhedonia, and reduces the ability to tolerate therapy, thereby contributing to increased mortality^25–27^. The causes and drivers of cachexia are currently not well understood but have been shown to involve a complex cross talk between the tumour and the host, involving several organs distant to the tumour such as the liver and brain^26–28^. While loss of appetite is a central feature of cachexia, it is not by itself sufficient to explain the syndrome. Several cytokine and hormonal mediators with pleiotropic roles signal to cachexia affected tissues such as the muscle, the brain and adipose tissue or the immune system have been identified. Tumour necrosis factor-α (TNFα), IL-6, Activin A and GDF15 have been implicated as cachexia drivers, although individually do not appear to mediate the full cachectic response^27–29^. For example, a humanized monoclonal antibody against GDF15 improved weight and physical activity in cancer patients with cachexia and elevated circulating GDF15 levels in a phase 2 clinical trial^30^. Importantly, cachexia cannot be reversed by conventional dietary approaches, such as supporting patients with increased nutrient and protein delivery^31^. Indeed, increased fat administration in certain preclinical cancer models, instead of replenishing adipose tissue, appeared detrimental and accelerated cachexia development^32^, while a ketogenic diet limited tumour growth but negatively affected the host^33^.

In light of the observed weight loss associated with the serine/glycine deficient diet, we sought to understand whether this diet impacts the onset or severity of cachexia. This concept is supported by a global multi-omics profiling study, which identified perturbations in one-carbon metabolism as a common modality across cachexia-targeted tissues, suggesting that serine metabolism may be involved in pathways linked to wasting, inflammation and energy regulation^34^. Surprisingly we found that – despite being lighter - mice maintained on the diet were more resistant to cachexia in several models of lung and pancreas cancer, even when the tumours were unresponsive to the diet. Mechanistically, serine/glycine dietary restriction reduced adiposity, which remained stable and was not further impacted by tumour presence, while enhancing hepatic antioxidant responses, decreasing lipid oxidation and attenuating systemic inflammatory signalling.

## RESULTS

### SGD diet decreases body weight and adiposity while preserving skeletal mass in WT C57Bl/6J mice

To investigate the systemic consequences of dietary serine and glycine restriction, wild-type (WT) C57Bl/6J mice were fed either a control or serine/glycine-deficient (SGD) diet. Male and female mice were age matched at the start of the intervention (Figure S1A, B) and body weight was monitored longitudinally (Figure 1A, B). SGD diet induced a rapid reduction in body weight in both sexes. Within the first week of dietary intervention, male mice lost significantly more weight than controls (- 6.62±2.37% versus -2.94±2.75%), while female mice displayed a similar but less pronounced response (-2.98±3.75% versus +0.87±2.44%) (Figure 1C, D). Despite this initial decline, SGD diet-fed mice subsequently maintained a stable body weight that was lower than control-fed animals throughout the study (Figure 1A,B & S1C,D).

**Figure 1.**
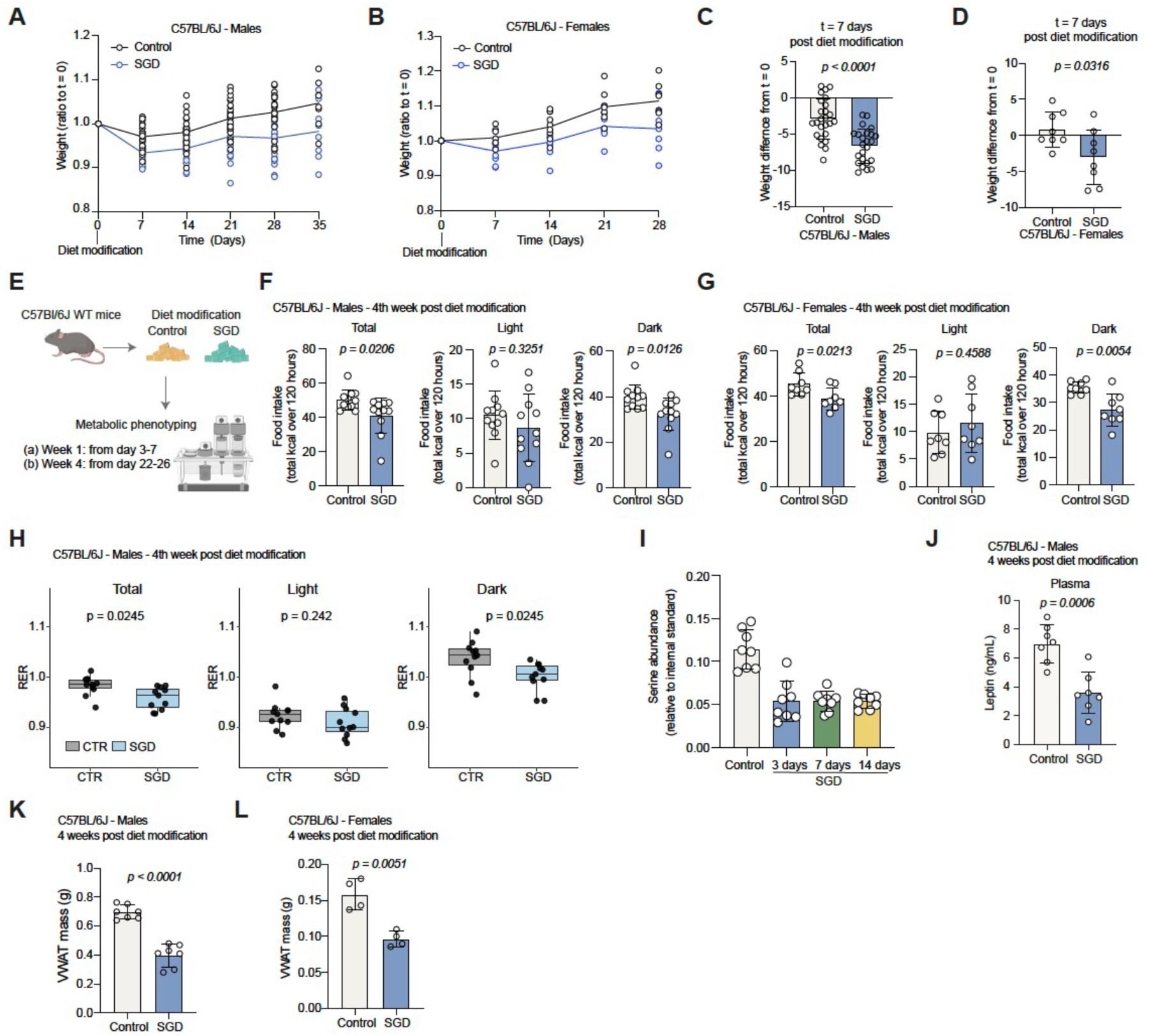
SGD diet decreases body weight and adiposity while preserving skeletal mass in WT C57Bl/6J mice. (A) Weight trajectories (ratio to t = 0) of C57Bl/6J WT male mice from diet enrolment till collection. Data are from n = 25 Control and 24 SGD mice. (B) Weight trajectories (ratio to t = 0) of C57Bl/6J WT female mice from diet enrolment till collection. Data are from n = 8 Control and 8 SGD mice. (C) Difference from starting weight of mice from (A) at t = 7 days. Values are mean±s.d. from n = 25 Control and 24 SGD mice. (D) Difference from starting weight of mice from (A) at t = 7 days. Values are mean±s.d. from n = 8 Control and 8 SGD mice. (E) Experimental strategy scheme for metabolic housing experiments. (F) Food intake (total kcal over 120 hours) for the total period, light phase and dark phase (as indicated) of C57Bl/6J WT male mice in control or SGD diet during the 4^th^ week post diet modification. Values are mean±s.d. from n = 11 control and n = 11 SGD mice. (G) Food intake (total kcal over 120 hours) for the total period, light phase and dark phase (as indicated) of C57Bl/6J WT female mice in control or SGD diet during the 4^th^ week post diet modification. Values are mean±s.d. from n = 8 control and n = 8 SGD mice. (H) Average RER of mice from (F) for the total period, light phase and dark phase (as indicated). Box-and-whisker plots showing median and interquartile range from n = 11 control and n = 11 SGD mice. (I) Serine levels in the circulation (plasma) of C57Bl/6J WT mice in control or SGD diet after 3, 7 and 14 days post diet modification. Values are mean±s.d.from n = 8 (4 female and 4 male) mice for each group (J) Plasma leptin levels of C57Bl/6J WT male mice in control or SGD diet 4 weeks post diet change. Values are mean±s.d. from n = 7 Control and 7 SGD mice. (K) VWAT mass (g) of C57Bl/6J WT male mice in control or SGD diet 4 weeks post diet modification. Values are mean±s.d.from n = 7 control and n = 7 SGD mice. (L) VWAT mass (g) of C57Bl/6J WT male mice in control or SGD diet 4 weeks post diet modification. Values are mean±s.d.from n = 7 control and n = 7 SGD mice. Statistical significance was assessed by two-tailed Welch’s t-test (C-D,F-G, J-L) and two-tailed unpaired t-test with Benjamini-Hochberg correction for multiple comparisons (H).

To characterise the metabolic adaptations underlying this phenotype, mice were housed in metabolic cages during the first week following dietary intervention and again after four weeks of feeding (Figure 1E). During the initial adaptation phase, SGD diet-fed male mice exhibited only a modest reduction in food intake (Figure S1E). In contrast, after four weeks of dietary restriction, both male and female SGD- fed mice consumed significantly less food than controls (Figure 1F,G). This reduction was primarily attributable to decreased food intake during the dark phase, when feeding activity is highest, whereas food consumption during the light phase was largely unchanged (Figure 1F,G). Importantly, ANCOVA analysis demonstrated that reduced food intake was independent of body mass, indicating that diet rather than body weight accounted for the observed differences in feeding behaviour (Figure S1F).

SGD administered male WT mice had a lower respiratory exchange ratio (RER) after 4 weeks of diet administration (Figure 1H), prominently in the dark phase of the cycle, maintaining their energy expenditure and ambulatory activity at similar levels as the controls (Figure S1G). However, female mice appeared to have similar RER and ambulatory activity, although they displayed lower energy expenditure (Figure S1H).

Targeted LCMS analysis confirmed that dietary restriction effectively depleted circulating serine. Plasma serine concentrations decreased markedly within three days of SGD feeding and remained suppressed thereafter, indicating rapid systemic adaptation to the diet (Figure 1I). Furthermore, SGD diet-fed mice had lower circulating leptin (Figure 1J).

Because SGD diet-fed mice maintained a leaner phenotype throughout the study, body composition was assessed at necropsy. Male mice exposed to SGD diet for four or six weeks exhibited a marked reduction in both visceral and subcutaneous white adipose tissue (WAT) depots (Figure 1K & S1I). In contrast, skeletal muscle mass was preserved, with no differences detected in gastrocnemius weights (Figure S1J). Consistent with maintained muscle integrity, SGD diet-fed mice retained normal grip strength throughout the intervention period (Figure S1K). Female mice similarly displayed reduced visceral adipose tissue mass after four weeks of SGD feeding (Figure 1L).

It has been previously reported that SGD diet reduces adiposity, without altering RER in WT animals^24^ thus adipose tissue loss may be the main consequence of SGD diet and differences in RER may be sex, age, strain or environment dependent. Collectively, our findings demonstrate that dietary serine and glycine restriction induces an acute reduction in body weight followed by long-term maintenance of a lean phenotype. This adaptation is characterised by progressive loss of adipose tissue, reduced food intake, while skeletal muscle mass and function remain preserved.

### SGD diet suppresses tumour progression and delays cachexia in Kras^G12D/+^;Lkb1^f/f^ NSCLC

Previous studies have demonstrated that dietary serine and glycine restriction can impair the growth of specific tumour types, including types of breast and colorectal cancer, although sensitivity varies considerably between cancer models^12–14,17,35,36^. To determine whether non-small cell lung cancers (NSCLC) are susceptible to serine/glycine restriction, we employed the Kras^G12D/+^;Lkb1^f/f^ (KL) genetically engineered mouse model, an aggressive NSCLC model characterised by rapid tumour progression and severe cancer-associated cachexia^37–39^.

Tumour formation was induced by intranasal delivery of AdCMV-Cre, and mice were switched to either control or SGD diet four weeks later (Figure 2A). SGD feeding significantly extended symptom-free survival, increasing median survival from 69 to 90 days post-Cre administration (Figure 2B). Longitudinal micro-computed tomography (μCT) imaging revealed that this survival benefit was associated with delayed tumour progression, as SGD-fed mice exhibited significantly slower expansion of lung lesions throughout the study (Figure 2C,D). Consistent with this observation, KL-derived tumour cells displayed reduced proliferation under serine/glycine-deprived conditions *in vitro* (Figure S2B) and tumours from SGD diet-fed mice contained fewer Ki67-positive cells at endpoint (Figure 2G,H). Despite these differences in tumour kinetics, mice from both dietary groups reached clinical endpoint with comparable tumour burden, as determined by lung mass and histopathological assessment (Figure 2E,F & S2A).

**Figure 2.**
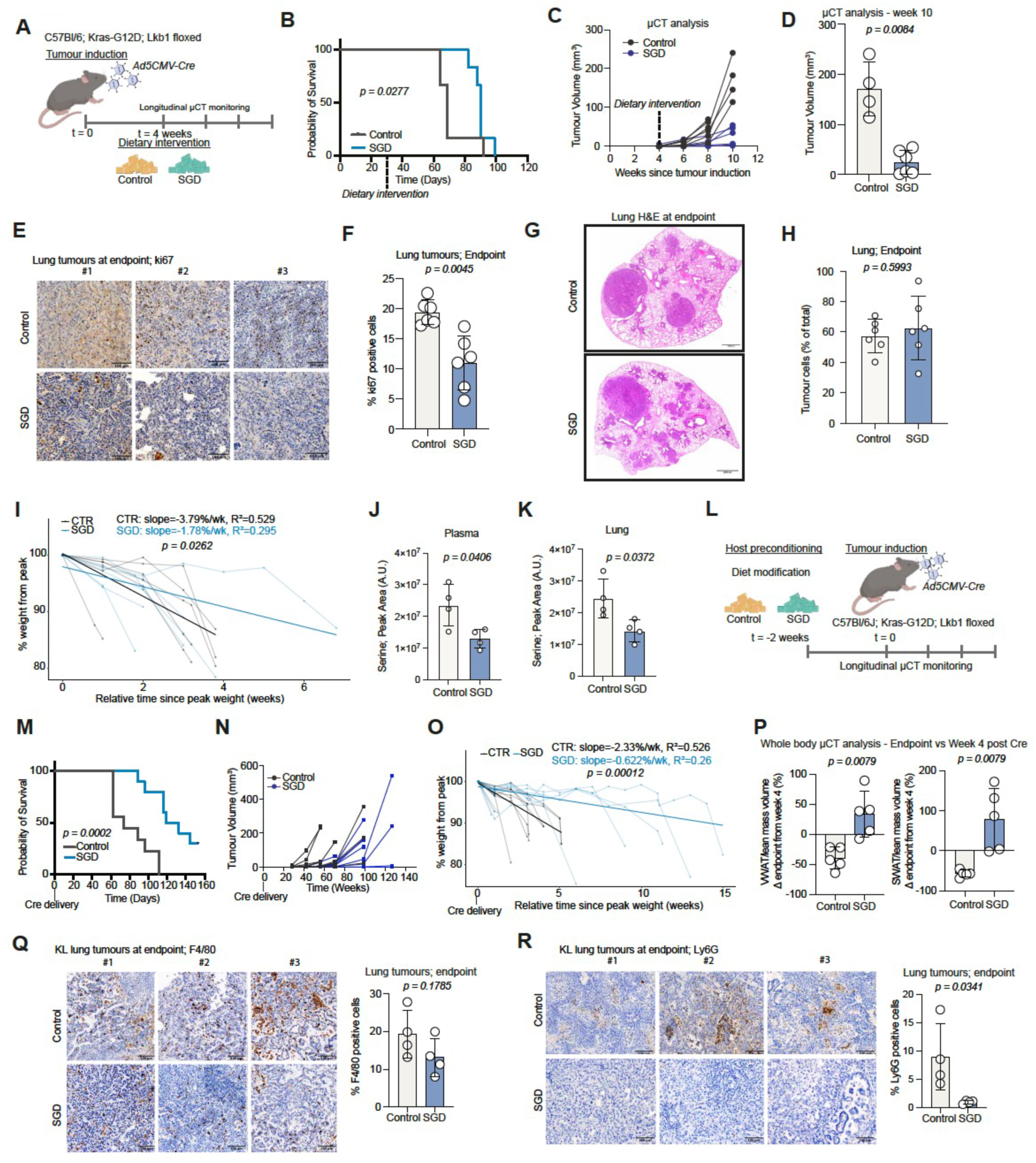
SGD diet suppresses tumour progression and delays cachexia in Kras^G12D/+^;Lkb1^f/f^ NSCLC. (A) Experimental strategy scheme of dietary intervention in Kras^G12D/+^;Lkb1^f/f^ mutant NSCLC (KL) model. For this cohort control diet was TestDiet, 5BQS and SGD TestDiet, 5BQT. (B) Probability of survival in KL male mice treated with control (n = 6) or SGD diet (n = 6). (C) Tumour volume (μCT quantification) trajectories in mice from (B). (D) Tumour volume (μCT quantification) at week 10 post Cre delivery. Data are from (C) shown as a bar plot with mean±s.d. values from n = 4 control and n = 6 SGD mice. (E) Representative pictures of ki67-stained lung tumours of mice from (B) at endpoint. Scale bars, 100 μm. Numbers indicate independent mice. (F) Quantification of ki67 positive nuclei of lung tumours from (F). Values are mean±s.d. of n = 6 control and n = 6 SGD mice. (G) Representative pictures of H&E-stained lungs of mice from (B) at endpoint. Scale bars, 1000 μm. (H) Quantification of tumour burden at endpoint in mice from (B,H). Values are mean±s.d. of n = 6 control and n = 6 SGD mice. (I) Weight loss trajectories from peak weight over time of mice from (B), with t = 0 corresponding to the timepoint of peak weight for each mouse. (J) Plasma serine levels of mice from (B) at endpoint. Values are mean±s.d. from n = 4 control and 4 SGD KL mice. (K) Lung tumour serine levels of mice from (B) at endpoint. Values are mean±s.d. from n = 4 control and 4 SGD KL mice. (L) Experimental strategy scheme of host dietary preconditioning in Kras^G12D/+^;Lkb1^f/f^ mutant NSCLC (KL) model. (M) Probability of survival in KL mice preconditioned with control (n = 9, 5 female & 4 male) or SGD diet (n = 10, 5 female and 5 male). (N) Tumour volume (μCT quantification) trajectories in mice from (M). (O) Weight loss trajectories from peak weight over time of mice from (M), with t = 0 corresponding to the timepoint of peak weight for each mouse. (P) Difference of VWAT (left) and SWAT (right) depots (μCT quantification) at endpoint compared to week 4, for mice with similar tumour burden at endpoint. Values are mean±s.d. from n = 5 control and 5 SGD mice. (Q) Representative pictures (left) of F4/80-stained lung tumours (as indicated) of mice from (B) at endpoint and quantification of F4/80 positive cells (right). Scale bars, 100 μm. Image numbers indicate independent mice. Values are mean±s.d. of n = 4 control and n = 4 SGD mice. (R) Representative pictures (left) of Ly6G-stained lung tumours (as indicated) of mice from (B) at endpoint and quantification of Ly6G positive cells (right). Scale bars, 100 μm. Image numbers indicate independent mice. Values are mean±s.d. of n = 4 control and n = 4 SGD mice. Statistical significance was assessed by two-tailed Mantel-Cox test (B,M), two-tailed Welch’s t-test (D,F,H,J,K,P) and two-tailed unpaired t-test (Q,R). Differences in trajectory slopes were assessed using a linear regression model with a group vs time interaction, evaluated by an F-test (I, O).

KL mice progressively develop cachexia during tumour progression and cachexia development in this model did not correlate with tumour burden^37,40–42^. Interestingly, SGD diet-fed mice exhibited a markedly slower decline in body weight compared with controls (Figure 2I & S2C). At endpoint, however, both groups displayed similarly reduced adipose tissue and skeletal muscle mass (Figure S2D,E), suggesting that SGD diet delayed rather than prevented tissue wasting. Serine concentrations remained lower than controls in both plasma and lung tissue of SGD-fed mice at endpoint (Figure 2J,K), whereas VWAT, quadriceps and liver were unaffected (Figure S2F). SGD KL mice at endpoint had similar circulating GDF15 levels (Figure S2G) and a trend for lower Activin A (Figure S2H), two reported drivers of cachexia in this model.

To determine whether prolonged adaptation to SGD diet could further influence disease progression, a second cohort was preconditioned with control or SGD diet for two weeks before tumour induction (Figure 2L). Under these conditions, the survival benefit was substantially enhanced, with median survival increasing from 74 to 125.5 days in SGD diet-fed animals (Figure 2M). As observed in the intervention cohort, tumour growth was significantly delayed (Figure 2N), whereas tumour burden at endpoint remained comparable between groups (Figure S2I,J). SGD-fed mice also experienced a more gradual decline in body weight throughout disease progression (Figure 2O and S2K). However, similarly to the intervention cohort, both groups displayed comparable levels of adipose tissue and skeletal muscle mass at endpoint (Figure S2L,M).

Because SGD diet-fed WT mice exhibited reduced adiposity, we next investigated adipose tissue dynamics during tumour progression using whole-body μCT imaging. Mice with comparable tumour burden at endpoint were selected for analysis (Figure S2N). At 4 weeks post Cre, when tumours were not detectable by μCT (Figure 2N), SGD diet-fed KL mice had lower VWAT and SWAT depots, as expected (Figure S2O). However, by endpoint, control-fed KL mice progressively lost both visceral and subcutaneous adipose tissue during tumour progression, while SGD diet-fed animals largely maintained these depots and, in some cases, exhibited modest increases over time (Figure 2P). Thus, despite reaching endpoint with similar adipose tissue mass, control and SGD diet-fed mice arrived at this state through fundamentally distinct trajectories: tumour-associated adipose wasting predominated in control animals, whereas SGD diet-fed mice underwent early dietary adaptation followed by relative preservation of adipose tissue during tumour progression.

Histological analysis further supported differences in adipose tissue remodelling between groups. Visceral adipose tissue from SGD diet-fed mice contained fewer F4/80-positive macrophages at endpoint (Figure S2P). Similarly, KL tumours from SGD diet-fed mice exhibited reduced Ly6G-positive neutrophil infiltration and a trend toward lower macrophage abundance (Figure 2R). Together, these findings demonstrate that SGD diet suppresses KL tumour growth while simultaneously delaying the onset of cachexia and attenuating inflammatory changes in both adipose tissue and the tumour microenvironment.

### SGD diet delays cachexia independently of tumour growth in Kras^G12D/+^;p53^f/f^ mutant NSCLC

To distinguish direct anti-tumour effects of SGD from host-mediated effects on cachexia, we next investigated the Kras^G12D/+^;p53^f/f^ (KP) NSCLC model, which displays slower disease progression and represent biologically distinct KRAS-driven NSCLC subtypes with differences in signalling, immune microenvironment and metabolic features^38^. KP tumours were induced by intranasal AdCMV-Cre delivery, and mice were switched to control or SGD diet six weeks later (Figure 3A). In contrast to the KL model where there was a clear benefit from the SGD diet with slower tumour growth and extended survival, SGD feeding did not affect overall survival, or tumour burden in KP mice, which reached endpoint at the same time as control fed mice (Figure 3B-D and S3A). At week 14, CT analysis suggested a trend towards slightly smaller tumours (S3B), although this was not significant and did not impact final tumour burden (Fig 3D). Likewise, KP-derived tumour cells proliferated normally in serine/glycine-depleted medium *in vitro* (Figure S3C) and tumour proliferation *in vivo* was unchanged, as assessed by Ki67 staining (Figure 3E,F). These findings indicate that KP tumours are intrinsically resistant to dietary serine and glycine restriction.

**Figure 3.**
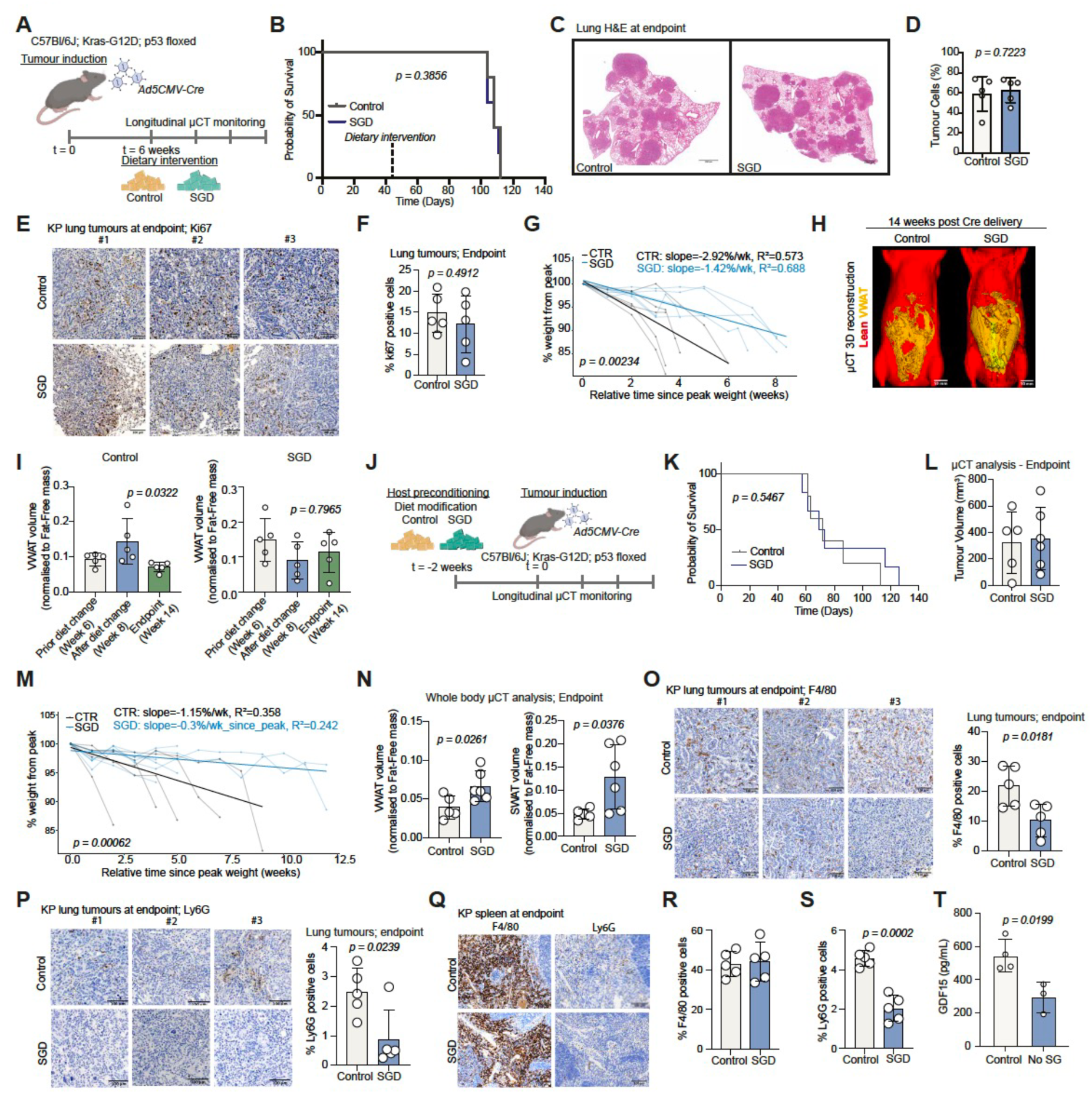
SGD diet delays cachexia independently of tumour growth in Kras^G12D/+^;p53^f/f^ mutant NSCLC. (A) Experimental strategy scheme of dietary intervention in Kras^G12D/+^;p53^f/f^ mutant NSCLC (KP) model. For this cohort control diet was TestDiet, 5BQS and SGD TestDiet, 5BQT. (B) Probability of survival in KP male mice treated with control (n = 5) or SGD diet (n = 5). (C) Representative pictures of H&E-stained lungs of mice from (B) at endpoint. Scale bars 1000 μm. (D) Quantification of tumour burden in mice from (B) at endpoint. Values are mean±s.d. of n = 5 control and 5 SGD mice. (E) Representative pictures of ki67-stained lung tumours of mice from (B) at endpoint. Scale bars, 100 μm. Numbers indicate independent mice. (F) Quantification of ki67 positive nuclei of lung tumours from (E). Values are mean±s.d. of n = 5 control and n = 5 SGD mice. (G) Weight loss trajectories from peak weight over time of mice from (B), with t = 0 corresponding to the timepoint of peak weight for each mouse. (H) Representative 3D rendering of lean mass and VWAT compartments generated from whole body μCT scans of control and SGD KP mice 14 weeks after Cre delivery. (I) VWAT volume (ratio to Fat-Free mass) of control (n = 5) (left panel) and SGD (n = 5) (right panel) KP mice before (week 6), after (week 8) diet modification and before endpoint. (J) Experimental strategy scheme of host dietary preconditioning in Kras^G12D/+^;p53^f/f^ mutant NSCLC (KP) model. (K) Probability of survival in KP mice treated with control (n = 5) or SGD diet (n = 6). (L) Tumour volume (μCT quantification) at endpoint in mice from (K). Values are mean±s.d. of n = 5 control and n = 6 SGD mice. (M) Weight loss trajectories from peak weight over time of mice from (K), with t = 0 corresponding to the timepoint of peak weight for each mouse. (N) VWAT (left panel) and SWAT (right panel) volume (ratio to Fat-Free mass) of control and SGD KP mice at endpoint. Values are mean±s.d. of n = 5 control and 6 SGD mice. (O) Representative pictures (left) of F4/80-stained lung tumours (as indicated) of mice from (B) at endpoint and quantification of F4/80 positive cells (right). Scale bars, 100 μm. Image numbers indicate independent mice. Values are mean±s.d. of n = 5 control and n = 5 SGD mice. (P) Representative pictures (left) of Ly6G-stained lung tumours (as indicated) of mice from (B) at endpoint and quantification of Ly6G positive cells (right). Scale bars, 100 μm. Image numbers indicate independent mice. Values are mean±s.d. of n = 5 control and n = 5 SGD mice. (Q) Representative pictures (left) of F4/80 and Ly6G-stained spleens of mice from (B) at endpoint. Scale bars, 100 μm. (R) Quantification of F4/80 positive cells from (Q). Values are mean±s.d. of n = 5 control and n = 5 SGD mice. (S) Quantification of Ly6G positive cells from (Q). Values are mean±s.d. of n = 5 control and n = 5 SGD mice. (T) Plasma GDF15 levels in mice from (B) at endpoint. Values are mean±s.d. of n = 4 control and 3 SGD mice. Statistical significance was assessed by two-tailed Mantel-Cox test (B,Κ), two-tailed Welch’s t-test (D,F,Ν,O,P,S,T) and ordinary one-way ANOVA with Tukey’s multiple comparisons test (I). Differences in trajectory slopes were assessed using a linear regression model with a group vs time interaction, evaluated by an F-test (G,Μ).

Despite the absence of an anti-tumour effect, SGD diet-fed KP mice exhibited significantly attenuated body weight loss during disease progression (Figure 3G and S3D). Endpoint adipose tissue and skeletal muscle masses were comparable between dietary groups (Figure S3E,F), suggesting that SGD diet primarily altered the kinetics of tissue wasting rather than its severity. We performed longitudinal μCT whole body analysis which revealed marked differences in body composition dynamics between the two groups. Control-fed mice progressively lost both visceral and subcutaneous adipose tissue between weeks 8 and 14 following tumour induction. In contrast, SGD diet-fed animals experienced an initial reduction in adipose tissue shortly after dietary intervention but subsequently maintained these depots throughout tumour progression (Figure 3H,I & S3G). Thus, as observed in KL mice, SGD diet-induced adipose tissue remodelling appeared to precede tumour development and protected against further tumour-associated adipose loss.

To determine whether extended adaptation to SGD diet altered disease outcome, a second cohort was preconditioned with diet for two weeks before tumour induction (Figure 3J). Consistent with the intervention study, SGD did not affect survival, tumour growth or endpoint tumour burden (Figure 3K,L and S3H). Nevertheless, SGD-fed mice displayed substantially improved preservation of body weight throughout disease progression (Figure 3M and S3I). Whereas control-fed mice reached endpoint following rapid body weight decline, SGD-fed animals maintained stable body weight and instead met endpoint criteria due to respiratory symptoms (rapid breathing) associated with advanced lung tumour burden.

Analysis of body composition at endpoint demonstrated significantly greater visceral and subcutaneous adipose tissue volumes in SGD-fed mice (Figure 3N). Consistently, these animals also showed greater visceral adipose tissue and gastrocnemius muscle mass at necropsy (Figure S3J,K). Visceral adipose tissue from SGD-fed mice also exhibited reduced macrophage infiltration (Figure S3L), suggesting diminished inflammatory remodelling.

Interestingly, despite their distinct responses to SGD diet in terms of tumour growth, KP tumours displayed reduced infiltration of F4/80-positive macrophages and Ly6G-positive neutrophils following dietary intervention, similar to KL tumours (Figure 3O,P). Furthermore, SGD-fed KP mice developed significantly smaller spleens than control animals in both experimental cohorts (Figure S3M). Although splenic macrophage abundance was unchanged (Figure 3Q,R), Ly6G-positive neutrophils were markedly decreased in spleens of SGD-fed animals (Figure 3Q,S). Consistent with these observations, circulating GDF15 concentrations were lower in SGD-fed KP mice at endpoint (Figure 3T).

Collectively, these findings demonstrate that SGD delays the onset of cachexia and the trajectory of weight loss in the KP model despite having no measurable effect on tumour growth. The preservation of adipose tissue and skeletal muscle, together with reduced systemic neutrophilia and lower circulating GDF15, suggests that dietary serine and glycine restriction modifies host responses to tumour burden independently of direct effects on tumour cell proliferation.

### SGD diet rewires hepatic metabolism and reduces systemic oxidative stress in KP NSCLC

Following the observation that SGD diet delays cachexia progression in KP mice, despite having no effect on tumour growth, we next sought to identify the host metabolic adaptations associated with this phenotype. We focused on the KP NSCLC model because tumour burden was comparable between dietary groups, allowing systemic effects of SGD diet to be investigated independently of differences in tumour progression.

To characterise tissue-specific metabolic responses, we performed polar metabolomics and lipidomics on liver, plasma, VWAT and gastrocnemius muscle collected from endpoint KP mice fed with either control or SGD diet (Figure 4A). Principal component analysis of untargeted polar metabolites revealed the clearest separation between dietary groups in the liver, whereas plasma, VWAT and skeletal muscle exhibited substantially weaker separation (Figure 4B and S4A), indicating that the liver is a major site of metabolic adaptation to serine and glycine restriction.

**Figure 4.**
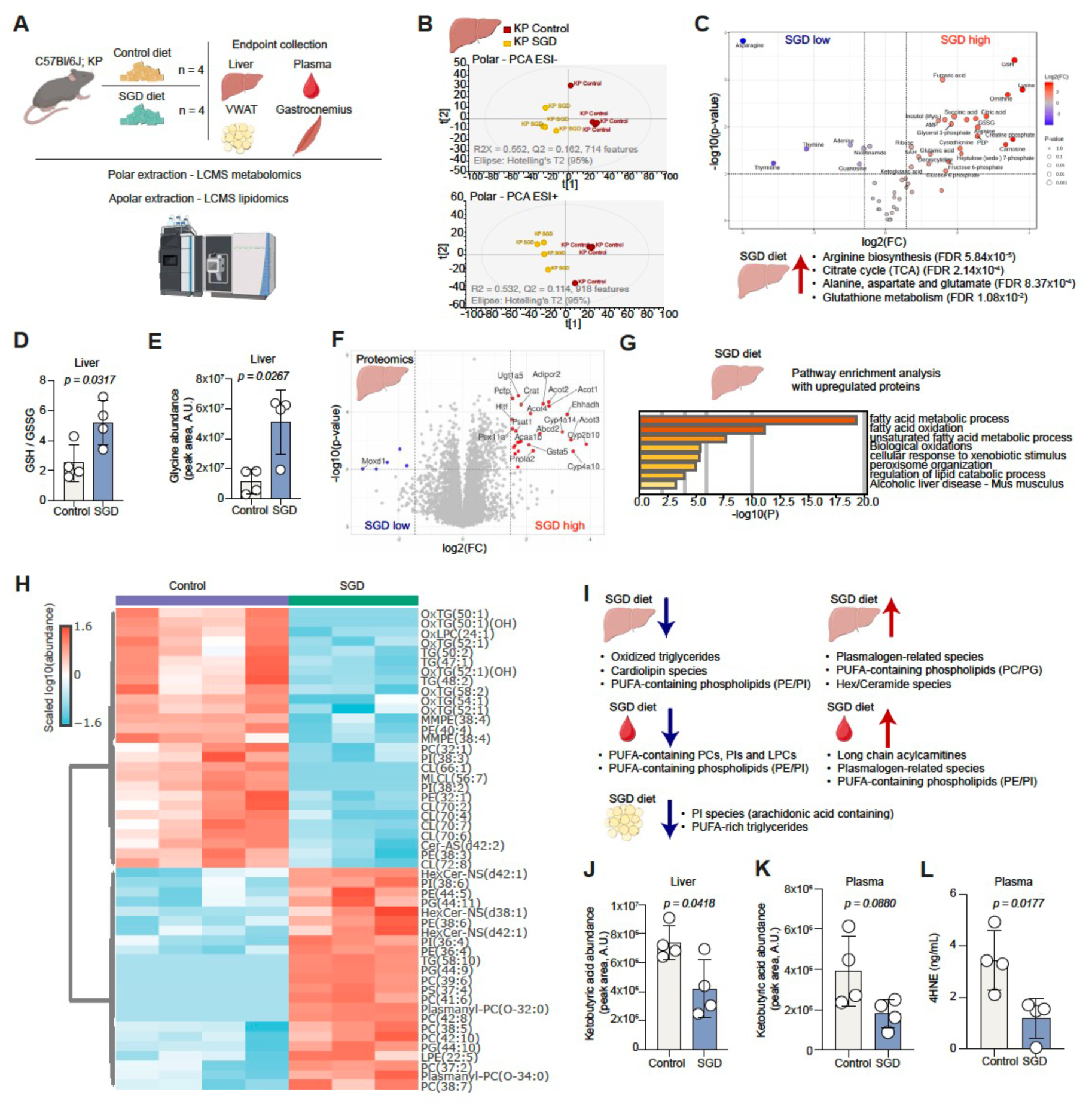
SGD diet rewires hepatic metabolism and reduces systemic oxidative stress in KP NSCLC. (A) Scheme depicting LCMS analysis strategy in tissues derived from the Kras^G12D/+^;p53^f/f^ mutant NSCLC (KP) model with control or SGD diet. (B) PCA plots of LCMS analysis of liver polar extracts from control or SGD-treated KP mice. (C) Volcano plot of identified metabolites of liver polar extracts from KP mice. Bottom; top enriched pathways from metabolites with increased abundance in SGD-treated mice. (D) GSH/GSSG ratio in liver lysates of KP mice treated with control or SGD diet. Values are mean±s.d. from n = 4 control and n = 4 SGD mice. (E) Glycine levels in liver lysates of KP mice treated with control or SGD diet. Values are mean±s.d. from n = 4 control and n = 4 SGD mice. (F) Volcano plot of detected proteins in liver lysates from control or SGD-treated KP mice. (G) Pathway enrichment analysis of upregulated proteins in liver lysates of SGD-treated mice from (F). (H) Heatmap, abundance of identified lipid species in apolar liver extracts from control or SGD treated KP mice. Only top 50 statistically significant identified lipid species are displayed. (I) Scheme depicting top categories of lipid species decreased (left) or increased (right) in liver (top), plasma (middle) and VWAT (bottom) extracts from KP mice treated with SGD diet. (J) Ketobutytic acid levels in liver lysates of KP mice treated with control or SGD diet. Values are mean±s.d. from n = 4 control and n = 4 SGD mice. (K) Ketobutytic acid levels in plasma of KP mice treated with control or SGD diet. Values are mean±s.d. from n = 4 control and n = 4 SGD mice. (L) Plasma 4HNE levels in KP mice treated with control or SGD diet at endpoint. Values are mean±s.d. of n = 4 control and 4 SGD mice. Statistical significance was assessed by two-tailed Welch’s t-test (D,E,J,K,L).

Targeted analysis of identified metabolites demonstrated a broad metabolic remodelling within the liver of SGD-fed mice (Figure 4C). Several tricarboxylic acid (TCA) cycle intermediates, including citrate, fumarate and succinate, were increased in abundance, together with metabolites associated with arginine metabolism. Notably, glutathione (GSH) levels were markedly elevated in SGD-fed livers, resulting in a significant increase in the GSH/GSSG ratio (Figure 4D and S4C,D). These findings suggest enhanced antioxidant buffering capacity in response to dietary amino acid restriction.

Despite sustained depletion of circulating serine, hepatic serine levels were maintained in SGD-fed mice, whereas glycine abundance was increased (Figure 4E and S4E,F). This pattern is consistent with compensatory adaptations within hepatic one-carbon metabolism that preserve intracellular serine availability during dietary restriction^43,44^ and provides glycine to support glutathione biosynthesis.

To investigate whether these metabolic changes were accompanied by broader alterations in hepatic function, we performed quantitative proteomic analysis of liver tissue. SGD feeding increased the abundance of multiple proteins involved in peroxisomal fatty acid metabolism and β-oxidation, including ACOT3, ACOT4, ABCD2, PEX11A and EHHADH (Figure 4F). Pathway enrichment analysis further revealed significant upregulation of fatty acid metabolic processes, lipid oxidation pathways and peroxisome-associated functions (Figure 4G), indicating extensive remodelling of hepatic lipid metabolism^45,46^.

To determine how these changes influenced hepatic lipid composition, we performed untargeted lipidomics. SGD-fed livers displayed substantial lipid remodelling characterised by reduced abundance of oxidised triglyceride species and ceramide-associated lipids, together with altered cardiolipin composition (Figure 4H and S4G,H). In contrast, plasmalogen-related phospholipids were enriched in SGD-fed animals, a lipid class closely linked to peroxisomal metabolism and frequently altered during oxidative and inflammatory stress^47–49^.

Similar signatures were detected systemically. Plasma lipidomics revealed increased abundance of long-chain acylcarnitines and plasmalogen-related species, together with reduced levels of several polyunsaturated phospholipids (Figure 4I and S4I-K). Plasmalogen phospholipids are generally considered to provide antioxidant function, in part by lowering polyunsaturated fatty acid (PUFA) oxidation and thereby limiting ROS-induced damage^50^. Consistent with a decrease in oxidative stress in the SGD fed mice, VWAT, exhibited lower abundance of PUFA-rich triglycerides and arachidonic acid-containing phosphatidylinositol species (Figure 4I and S4L).

To assess whether enhanced lipid utilisation was associated with increased ketogenesis, we quantified ketobutyric acid levels in liver and plasma. Surprisingly, ketobutyric acid abundance was reduced in both compartments following SGD diet (Figure 4J,K), indicating that the observed increase in fatty acid metabolic pathways does not result in accumulation of ketone-related intermediates.

Because lipid oxidation can promote oxidative stress through the generation of reactive lipid species, we next measured circulating concentrations of 4-hydroxynonenal (4-HNE), a major lipid peroxidation product derived from polyunsaturated fatty acids^51^. SGD-fed mice exhibited significantly lower plasma 4-HNE levels than controls (Figure 4L), reflecting reduced systemic oxidative stress and consistent with the changes in lipids noted above.

Collectively, these findings identify a coordinated hepatic adaptation to dietary serine and glycine restriction characterised by enhanced antioxidant capacity, activation of peroxisomal lipid metabolism and extensive lipid remodelling. These changes are accompanied by reduced systemic lipid peroxidation and may contribute to the delayed cachexia phenotype observed in SGD-fed tumour-bearing mice.

### SGD diet improves host responses in SGD-resistant models of PDAC

To determine whether SGD diet influences the host in other tumour types, we investigated pancreatic ductal adenocarcinoma (PDAC) driven by Kras^G12D^ and p53^R172H^ mutations (KPC). Previous studies have reported that KPC tumours are largely resistant to dietary serine and glycine restriction^13^. We therefore asked whether SGD could nevertheless influence the host response to tumour burden independently of effects on tumour growth.

To minimise variability in tumour initiation and to allow collection at an early stage of systemic disease, KPC cells were orthotopically transplanted into the pancreas of C57BL/6J mice that had been preconditioned with either control or SGD diet (Figure 5A). Mice were analysed three weeks after transplantation, when pancreatic tumours were established but before the development of advanced disease. Consistent with previous reports, KPC cells proliferated normally under serine/glycine-deprived conditions in vitro (Figure S5A), and tumour mass at collection was comparable between dietary groups (Figure S5B), confirming resistance to SGD diet.

**Figure 5.**
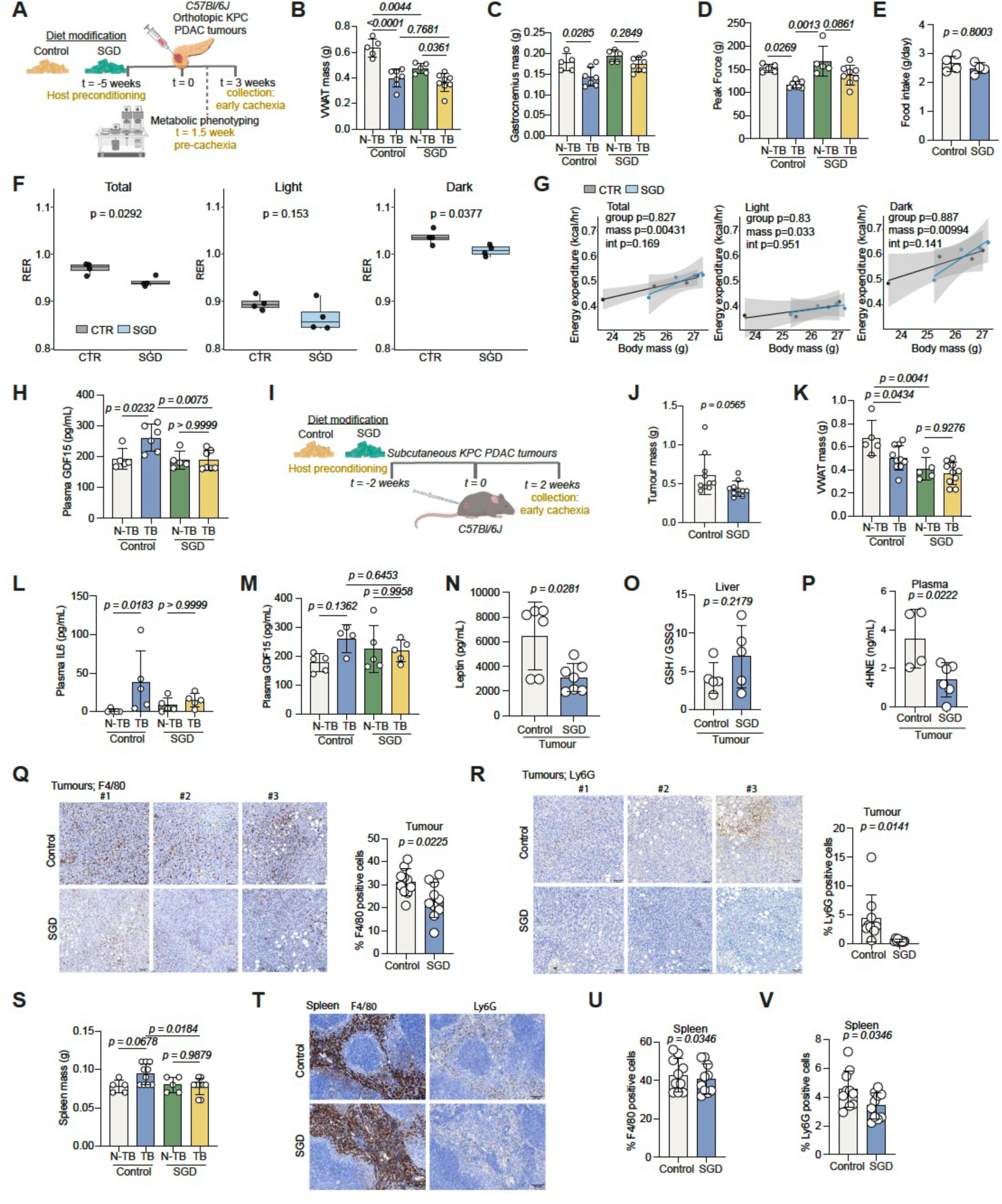
SGD diet improves host responses in SGD-resistant models of PDAC. (A) Experimental strategy scheme of host dietary preconditioning in Kras^G12D/+^;p53^R172H/+^ mutant (KPC) orthotopic PDAC model. (B) VWAT mass of mice from (A). Values are mean±s.d. from n = 5 WT control diet, n = 7 tumour control diet, n = 5 WT SGD diet and n = 8 tumour SGD diet. (C) Gastrocnemius mass of mice from (A). Values are mean±s.d. from n = 5 WT control diet, n = 7 tumour control diet, n = 5 WT SGD diet and n = 8 tumour SGD diet. (D) Peak force (g) of of mice from (A). Values are mean±s.d. from n = 5 WT control diet, n = 8 tumour control diet, n = 5 WT SGD diet and n = 8 tumour SGD diet. (E) Food intake (g/day) of of mice from (A). Values are mean±s.d. from n = 4 control and n = 4 SGD diet mice. (F) Average RER of mice from (A) for the total period, light phase and dark phase (as indicated). Box-and-whisker plots showing median and interquartile range from n = 11 control and n = 11 SGD mice. (G) Energy expenditure (kcal/hr) over body mass (g) of mice from (A). Values are from n = 4 Control and 4 SGD mice for the total period, light phase and dark phase (as indicated). Solid lines represent fitted values from ANCOVA analysis with group as a fixed factor and body mass as a covariate; shaded regions denote 95% confidence intervals. (H) Plasma GDF15 levels of mice from (A). Values are mean±s.d. from n = 5 WT control diet, n = 6 tumour control diet, n = 5 WT SGD diet and n = 7 tumour SGD diet. (I) Experimental strategy scheme of host dietary preconditioning in Kras^G12D/+^;p53^R172H/+^ mutant (KPC) subcutaneous PDAC model. (J) Tumour mass (g) at endpoint in mice from (I). Values are mean±s.d. from n = 10 control and 10 SGD mice. (K) VWAT mass in mice from (I). Values are mean±s.d. from n = 5 WT control diet, n = 10 tumour control diet, n = 5 WT SGD diet and n = 10 tumour SGD diet. (L) Plasma IL-6 levels in mice from (I). Values are mean±s.d. from n = 5 WT control diet, n = 5 tumour control diet, n = 5 WT SGD diet and n = 5 tumour SGD diet. (M) Plasma GDF15 levels in mice from (I). Values are mean±s.d. from n = 5 WT control diet, n = 4 tumour control diet, n = 5 WT SGD diet and n = 4 tumour SGD diet. (N) Plasma leptin levels in mice from (I). Values are mean±s.d. from n = 6 tumour control diet and n = 7 tumour SGD diet. (O) GSH/GSSG ratio in liver lysates of mice from (I). Values are mean±s.d. from n = 5 control and n = 5 SGD mice. (P) 4HNE levels in plasma of mice from (I). Values are mean±s.d. from n = 4 control and n = 6 SGD mice. (Q) Representative pictures (left) of F4/80-stained KPC subcutaneous tumours (as indicated) of mice from (I) and quantification of F4/80 positive cells (right). Scale bars, 100 μm. Image numbers indicate independent mice. Values are mean±s.d. of n = 5 WT control diet, n = 10 tumour control diet, n = 5 WT SGD diet and n = 10 tumour SGD diet. (R) Representative pictures (left) of Ly6G-stained KPC subcutaneous tumours (as indicated) of mice from (I) and quantification of Ly6G positive cells (right). Scale bars, 100 μm. Image numbers indicate independent mice. Values are mean±s.d. of n = 5 WT control diet, n = 10 tumour control diet, n = 5 WT SGD diet and n = 10 tumour SGD diet. (S) Spleen mass in mice from (I). Values are mean±s.d. from n = 5 WT control diet, n = 10 tumour control diet, n = 5 WT SGD diet and n = 10 tumour SGD diet. (T) Representative pictures of F4/80 and Ly6G-stained spleens of mice from (I) Scale bars, 100 μm. Image numbers indicate independent mice. (U) Quantification of F4/80 positive cells from (T). Values are mean±s.d. of n = 10 tumour control diet and n = 10 tumour SGD diet. (V) Quantification of Ly6G positive cells from (T). Values are mean±s.d. of n = 10 tumour control diet and n = 10 tumour SGD diet. Statistical significance was assessed by ordinary one-way ANOVA with Tukey’s multiple comparisons test (B,C,H,K,M,S), by ordinary one-way ANOVA with Šidák’s multiple comparisons test (D), by two tailed unpaired Welch’s t-test (E,J,N,O,Q,R,U,V), two-tailed unpaired t-test (P), Krusjal-Wallis with Dunn’s multiple comparisons test (L) and two-tailed unpaired t-test with Benjamini-Hochberg correction for multiple comparisons (F).

Despite similar tumour burden, tumour-bearing mice fed with the control diet displayed greater loss of visceral adipose tissue than SGD-fed animals (Figure 5B). Skeletal muscle mass was also improved in the SGD group (Figure 5C). Consistent with these changes, grip strength measurements demonstrated impaired muscle function in tumour-bearing mice fed the control diet, whereas SGD-fed tumour-bearing animals were comparable to non-tumour controls (Figure 5D).

To determine whether these differences were associated with altered energy balance, mice were housed in metabolic cages beginning 1.5 weeks after tumour implantation. Food intake was similar between tumour-bearing mice fed control or SGD diets (Figure 5E), suggesting that tumour presence may have a greater impact on the control fed animals in terms of appetite. As observed in healthy mice, SGD-fed animals exhibited a lower RER (Figure 5F and S5C). As expected, energy expenditure correlated with body mass, but mass-adjusted energy expenditure and ambulatory activity were similar between groups (Figure 5G & S5C). Although both groups displayed mild body weight loss relative to pre-surgical values (Figure S5D), circulating GDF15 concentrations were elevated only in tumour-bearing mice fed the control diet and remained low in SGD diet-fed animals (Figure 5H).

To further investigate early systemic manifestations of tumour burden in a model not susceptible to local pancreatic invasion and metastatic spread, which often occurs in the orthotopic model, KPC cells were implanted subcutaneously into mice preconditioned with either control or SGD diet (Figure 5I). As in the orthotopic model, there was no significant change in terms of tumour growth by SGD diet feeding (Figure 5J). At this early stage of disease, overall body weight changes were modest (Figure S5E), and no significant skeletal muscle wasting was evident (Figure S5F,G). However, control-fed tumour-bearing mice had already begun to lose visceral adipose tissue, whereas SGD diet-fed mice maintained adipose stores (Figure 5K). Control-fed tumour-bearing animals also exhibited elevated circulating IL-6 concentrations, whereas SGD-fed mice did not (Figure 5L). In contrast, circulating GDF15 levels remained similar between groups at this stage (Figure 5M). As expected from the lower overall adiposity of SGD-fed mice, independent of tumour presence, circulating leptin concentrations were reduced relative to controls (Figure 5N).

We next investigated whether the metabolic adaptations identified in KP mice were also present in tumour-bearing KPC hosts. Although animals were collected at an earlier stage of disease progression, SGD-fed mice displayed a trend towards increased hepatic and circulating GSH/GSSG ratios (Figure 5O and S5H), suggesting enhanced antioxidant capacity. Consistent with observations in KP mice, plasma concentrations of 4-HNE were significantly reduced in SGD-fed animals (Figure 5P), indicating lower systemic oxidative stress.

Analysis of the tumour microenvironment revealed reduced infiltration of both F4/80-positive macrophages and Ly6G-positive neutrophils in KPC tumours from SGD-fed mice (Figure 5Q,R). Furthermore, tumour-bearing mice fed the control diet had increased spleen mass, whereas this was prevented by SGD feeding (Figure 5S). Although splenic macrophage abundance was unchanged between groups (Figure 5T,U), SGD-fed mice exhibited significantly fewer Ly6G-positive neutrophils within the spleen (Figure 5T,V).

Together, these findings demonstrate that dietary serine and glycine restriction improves host responses to PDAC, despite having no effect on tumour growth. Preservation of adipose tissue and muscle function, reduced systemic oxidative stress, diminished splenic and systemic neutrophil accumulation and attenuation of inflammatory signalling suggest that SGD induces a host-protective programme.

### SGD diet suppresses acute systemic inflammatory signalling associated with cachexia

Having observed reduced neutrophil accumulation, lower circulating inflammatory mediators and delayed cachexia progression in both KP NSCLC and KPC PDAC models, we next sought to investigate the broader inflammatory consequences of dietary serine and glycine restriction. To identify signalling pathways consistently altered by SGD, we performed cytokine array profiling on plasma and liver lysates collected from tumour-bearing KP NSCLC and KPC PDAC mice fed either control or SGD diet (Figure 6A). Comparative analysis revealed a striking suppression of inflammatory signalling across both tumour models. In plasma, five cytokines were consistently increased whereas sixteen cytokines were commonly decreased in SGD diet-fed mice (Figure S6A-C). In liver lysates, twenty cytokines were commonly lower by SGD diet feeding (Figure S6D,E), suggesting that dietary serine and glycine restriction broadly dampens systemic inflammatory activation.

**Figure 6.**
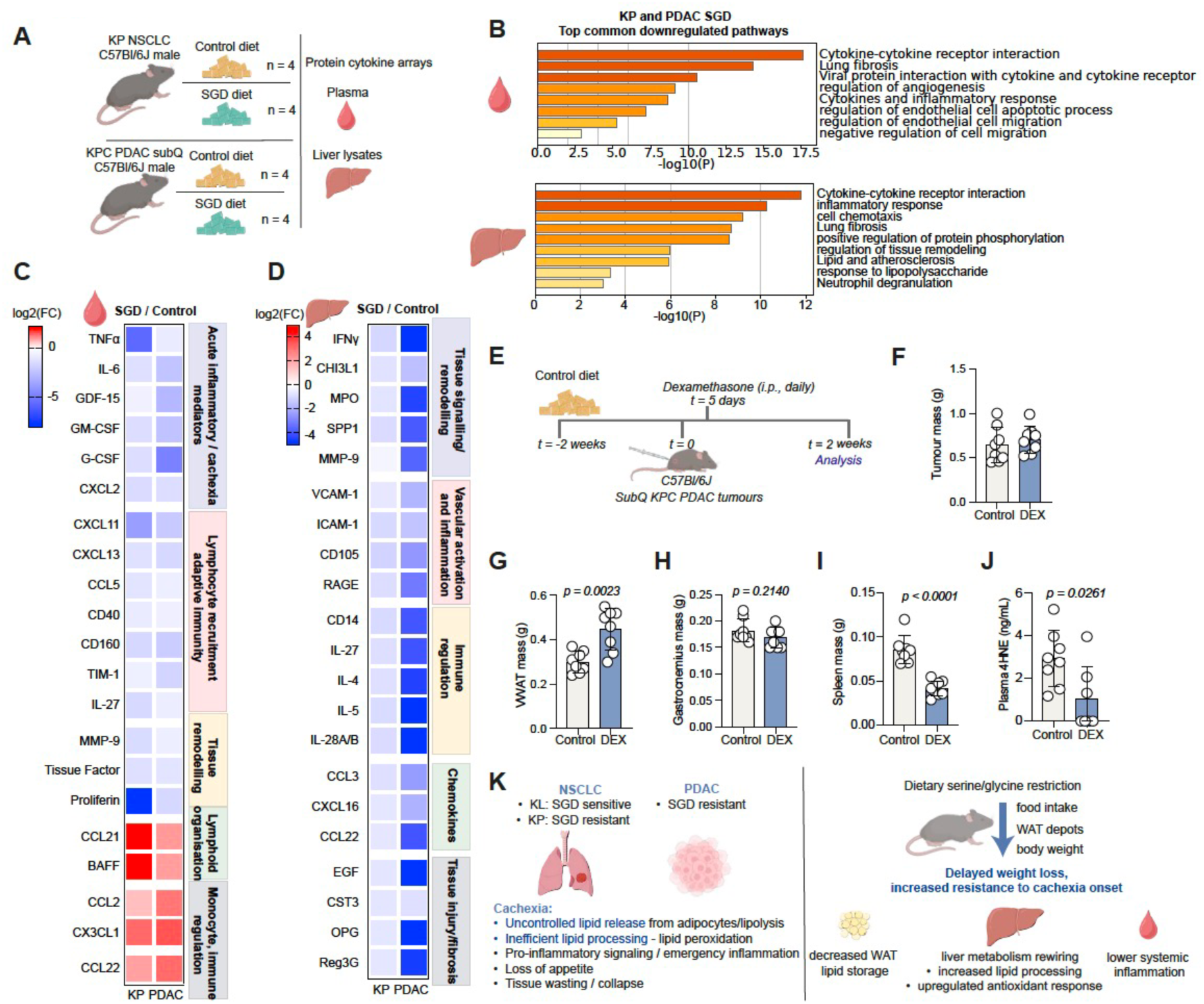
SGD diet suppresses acute systemic inflammatory signalling associated with cachexia. (A) Scheme depicting cytokine analysis strategy in tissues derived from the Kras^G12D/+^;p53^f/f^ mutant NSCLC (KP) and the Kras^G12D/+^;p53^R172H/+^ mutant (KPC) subcutaneous PDAC.models with control or SGD diet. (B) Pathway enrichment analysis of common downregulated cytokines in SGD-treated KP and PDAC models from (A). Top; plasma. Bottom; liver lysates. (C) Heatmap, plasma cytokines commonly down- and up-regulated in SGD-treated KP and PDAC models from (A). (D) Heatmap, liver cytokines commonly downregulated in SGD-treated KP and PDAC models from (A). (E) Experimental strategy scheme of dexamethasone administration in Kras^G12D/+^;p53^R172H/+^ mutant (KPC) subcutaneous PDAC mice. (F) Tumour mass (g) at collection in mice from (E). Values are mean±s.d. from n = 8 control and 8 DEX mice. (G) VWAT mass (g) at collection in mice from (E). Values are mean±s.d. from n = 8 control and 8 DEX mice. (H) Gastrocnemius mass (g) at collection in mice from (E). Values are mean±s.d. from n = 8 control and 8 DEX mice. (I) Spleen mass (g) at collection in mice from (E). Values are mean±s.d. from n = 8 control and 8 DEX mice. (J) Plasma 4HNE in mice from (E). Values are mean±s.d. from n = 8 control and 7 DEX mice. (K) Schematic summary. Statistical significance was assessed by two tailed unpaired Welch’s t-test (G,H,I,J).

Pathway enrichment analysis of commonly downregulated cytokines revealed significant suppression of pathways associated with inflammatory signalling, cytokine-receptor interactions, tissue remodelling, fibrosis and immune cell recruitment in both plasma and liver (Figure 6B). These observations are consistent with the reduced inflammatory phenotype observed across multiple tumour models. Among the most prominently reduced circulating factors were several mediators previously implicated in cachexia progression, including TNFα, IL-6 and GDF15, together with the myelopoietic factors G-CSF and GM-CSF, which contribute to systemic inflammation, neutrophil expansion and cachexia-associated host remodelling^27^ (Figure 6C). In addition, SGD feeding reduced multiple chemokines involved in immune cell recruitment, including CXCL2, CXCL11, CXCL13 and CCL5, as well as factors associated with tissue remodelling and vascular activation, such as MMP9 and Tissue Factor (Figure 6C). Analysis of liver lysates identified a similar pattern, with decreased abundance of inflammatory cytokines, chemokines and tissue-remodelling factors including IFNγ, SPP1, MPO, VCAM1, ICAM1, CCL3 and CXCL16 (Figure 6D). Collectively, these data indicate that SGD induces a broad suppression of inflammatory and tissue-remodelling programmes associated with cancer progression and cachexia.

To determine whether limiting inflammation could contribute directly to the protective effects observed in SGD diet-fed mice, we treated mice bearing subcutaneous KPC tumours with dexamethasone, a potent anti-inflammatory glucocorticoid, while maintaining animals on a control diet (Figure 6E). Dexamethasone did not alter tumour mass at endpoint (Figure 6F), allowing host responses to be evaluated independently of changes in tumour burden. Remarkably, dexamethasone-treated mice retained significantly more visceral adipose tissue than vehicle-treated controls (Figure 6G). Gastrocnemius muscle mass was unchanged between groups (Figure 6H), although this model does not exhibit substantial muscle wasting at the selected collection time point. Dexamethasone treatment also markedly reduced spleen mass (Figure 6I), consistent with suppression of systemic inflammatory activation and emergency myelopoiesis.

Analysis of circulating cytokines demonstrated that dexamethasone broadly decreased many of the same inflammatory mediators that were suppressed by SGD feeding (Figure S6F), suggesting convergence on overlapping inflammatory pathways. Furthermore, dexamethasone-treated animals exhibited lower plasma concentrations of the lipid peroxidation product 4-HNE (Figure 6J), indicating that suppression of inflammatory signalling is associated with reduced systemic oxidative stress.

Cancer cachexia is characterised by progressive anorexia, adipose and skeletal tissue wasting, chronic inflammation and metabolic dysfunction. Across multiple tumour models, dietary serine and glycine restriction consistently delayed the onset of these pathological features despite variable effects on tumour growth. Overall, our data suggest that SGD diet induces a systemic metabolic adaptation in the host, characterised by reduced adipose tissue storage, enhanced hepatic lipid processing, increased antioxidant capacity and suppression of inflammatory signalling (Figure 6K). These adaptations limit the accumulation of lipid peroxidation products, limit neutrophil-associated inflammatory responses and ultimately increase resistance to cachexia development. Thus, dietary serine and glycine restriction exerts beneficial effects on the host that extend beyond direct regulation of tumour growth and may represent a strategy to improve resilience to cancer-associated systemic disease.

## DISCUSSION

Dietary serine/glycine restriction results in a modest reduction in body weight in mice, which stabilizes over time and is not associated with overt impairment of health and wellbeing. Indeed, previous studies have reported maintaining mice on the diet for up to a year, with minimal detectable impact on health^14^. In the present study, we observed that serine/glycine restriction induces a limited reduction in adiposity, without affecting muscle mass or grip strength. The reduction in body weight appeared at least partly attributable to decreased food intake and may therefore reflect features of mild caloric restriction, although metabolic adaptations likely contribute as well.

Strikingly, in multiple pancreatic and lung cancer models, serine/glycine restricted mice exhibited delayed onset and development of cachexia and a relative preservation of adipose tissue compared to control-fed animals. Importantly, this effect was evident even in models of cancer where the tumour did not respond to the diet, showing that protection from cachexia was not secondary to reduced tumour burden but instead reflects a host-mediated adaptation.

Metabolomic analysis showed a clear reprogramming of liver metabolism, especially with respect to lipid handling. While the overall change in lipid metabolism in diet fed mice was complex, a clear reduction in oxidized lipid species was observed alongside decreased circulating 4-hydroxynonenal (4-HNE), indicating less systemic lipid peroxidation and oxidative stress. The whole-body response of reduced oxidative damage is an interesting counterpoint to the increase in oxidative stress seen in tumour cells deprived of serine and glycine – likely reflecting a decrease in glutathione availability. The systemic response was accompanied by reduced inflammatory signalling and diminished neutrophil infiltration into tumours, consistent with a broadly attenuated inflammatory state. Given that cachexia is strongly associated with systemic inflammation, including TNF, IL-6 and IFNγ signalling from tumour, stromal and immune compartments, this reduction in inflammatory status is likely to contribute to the delayed wasting phenotype observed. These cytokines are known to directly promote adipose and muscle catabolism through transcriptional reprogramming of metabolic and inflammatory pathways in peripheral tissues^27–29^.

Cachexia is further driven by endocrine mediators such as GDF15, which acts centrally to suppress appetite and increase lipid mobilisation^29^. While anorexia contributes variably across tumour types, it is not sufficient to explain cachexia in all settings, and in several models, cachexia develops independently of reduced food intake^25,52,53^. In this context, the reduced adiposity induced by serine/glycine restriction suggests that baseline metabolic state influences susceptibility to cachectic progression.

Mechanistically, adipose tissue represents an early and central node in cachexia progression, with adipocyte lipolysis increasing circulating fatty acids and glycerol that fuel systemic metabolic reprogramming^29,54,55^. Consistent with this, experimental inhibition of lipolysis can attenuate features of cachexia even in the presence of ongoing inflammation^56–58^. Our findings suggest that serine/glycine restriction may induce a metabolically distinct adipose state that is less susceptible to tumour-induced lipolytic signalling and increased inflammation.

Interestingly we observed that serine/glycine restriction improves hepatic peroxisomal lipid handling and reduces lipid peroxidation without significantly altering hepatic ketogenesis. This distinction is important in light of previous work showing that restoration of ketone production via PPARα activation can ameliorate cachexia-like phenotypes in certain genetic models ^37^. However, increasing circulating ketones through dietary ketogenic interventions alone has not consistently improved cachexia outcomes or survival in preclinical studies^59^. Our data are therefore consistent with the interpretation that improved peroxisomal lipid detoxification and reduced oxidative lipid stress, rather than ketone availability *per se*, may be more relevant for systemic protection against cachexia in this context. This suggests that replacing metabolic outputs, such as ketones, will be insufficient if upstream lipid peroxidation and inflammatory lipid signalling are not resolved.

While serine restriction has been reported to impair muscle regeneration and reduce muscle stem cell function in injury models ^60^, as well as to decrease myotube width *in vitro* and induce muscle wasting in subcutaneous tumour implantation models of colorectal cancer cells in Balb/c and Foxn1 nu/nu mice^61^, we did not observe reductions in muscle mass or function in tumour-bearing animals under our experimental conditions. These findings indicate that the impact of serine restriction on skeletal muscle is context-dependent, varying with tumour type, systemic metabolic state and duration of dietary intervention. Importantly, early reductions in body weight following dietary intervention primarily reflect dietary acclimatisation and are not indicative of tumour-driven wasting.

Together, our findings support a model in which serine/glycine restriction induces a metabolically reprogrammed host state characterised by reduced oxidative lipid stress, improved peroxisomal lipid handling and dampened systemic inflammation. This environment appears to delay activation of cachexia-associated pathways, thereby preserving adipose tissue despite ongoing tumour growth. Importantly, these effects are observed independently of tumour sensitivity to serine restriction.

The mechanisms linking serine/glycine restriction to reduced inflammation remain incompletely understood. One possibility is that altered redox balance and lipid handling decrease inflammatory signalling in cachexia-affected tissues. Changes in adipose tissue volume, function and remodelling may also contribute to altered cytokine production in the host. In addition, serine absence may also impact haematopoietic responses, including pathways involved in emergency granulopoiesis and neutrophil production. Our findings are further supported by the accompanying study by Gomez-Jenkins et al. showing a similar delay in cachexia in mice subjected to caloric restriction, further highlighting the proposal that the baseline metabolic state of the host is closely linked to inflammatory status and systemic response during tumorigenesis. This study shows that blocking a single cytokine is not sufficient to prevent cachexia, suggesting that intervention at multiple points will be required. Instead, caloric restriction preconditioning enforced a normal host protective response to nutrient limitation, attenuating cachexia through decreased multi-tissue inflammation and catabolism. The present work and that from Gomez-Jenkins et al indicate that the broad programme of metabolic, antioxidant and inflammatory changes elicited by either the serine/glycine restricted or the caloric restricted diet may be key to successful control of cachexia. These studies also suggest that - contrary to the dogma that increased calories must be offered to counteract cachexia - dietary nutrient limitation, either through amino acid restriction or caloric restriction, elicits a host response that limits cachexia.

In NSCLC tumours driven by Kras and Lkb1 mutations, but not by Kras and p53, both the tumours and the host responded to dietary serine/glycine restriction. Further work will be required to disentangle the relative contributions of direct effects on the tumor, adipose tissue remodelling, hepatic lipid detoxification and immune modulation in mediating this protective phenotype. Nonetheless, our data suggest that serine/glycine restriction will have broader therapeutic implications beyond direct tumour targeting, potentially improving host metabolic resilience even in cancers that are not metabolically responsive to this intervention.

## Acknowledgments

We thank the following: Julian Downward and Christopher Moore for breeding and providing Kras^G12D^;Lkb1^fl^ (KL) and Kras^G12D^;p53^fl^ (KP) mice as well as the KL cell line, Julia Weber for providing Kras^G12D^;p53^fl^ (KP) mice, the Francis Crick Institute’s Proteomics STP and especially Christelle Dubois for proteomics experiments, the Metabolomics, Cell Science and Experimental Histopathology STPs, specifically Thomas Snoeks and the In Vivo Imaging STP for guidance with *in vivo* lung and whole body μCT imaging and analysis. We are grateful to the Francis Crick Institute’s Biological Research Facility (BRF) for their excellent support of *in vivo* experiments and in particular Eniko Kiraly Avci and Rekha Subramaniam for their contribution in establishing an indirect calorimetry system. We also thank the BRF surgical services team, namely Sophie Wood, Sarah Phelan and the BRF veterinarian Arturo Fernandez, for their generous support with orthotopic pancreas surgeries and the pathologist Alejandro Suarez-Bonnet for independently quantifying and scoring lung tumour burden in NSCLC models. K.H.V. is funded by ERC-2020-ADG PancObese 10102064 and supported by the Francis Crick Institute which receives its core funding from Cancer Research UK (CC2073), the UK Medical Research Council (CC2073) and the Wellcome Trust (CC2073). V.P. has a Cancer Research UK Grant Award for Cancer Grand Challenges CANcer Cachexia Action Network (CANCAN) (PRJ_20994). We are also grateful to the Cancer Grand Challenge CANCAN consortium for helpful discussion and advice.

## Author contributions

V.P. and K.H.V. designed the research; V.P. performed research; J.C. carried out histopathological analyses; M.M.Z. carried out dissections and helped with mouse surgeries; I.F. performed LC-MS/GC-MS; V.P. and K.H.V. analysed data; and V.P. and K.H.V. wrote the paper.

## MATERIALS AND METHODS – STAR METHODS

### EXPERIMENTAL MODELS AND STUDY DETAILS

All *in vivo* experiments were conducted in compliance with the with the Animals (Scientific Procedures) Act 1986 and the EU Directive 2010 (PPL PP1389725) and within institutional welfare and ethical guidelines of the Francis Crick Institute (establishment licence X49D044E1). Mice had *ad libitum* access to food and water and were kept in a 12h day/night cycle (7:00 to 19:00) in rooms at 21°C and 55% humidity. Mice were acclimatized to their environment for at least one week prior to experimentation. Mice used in the experiments were randomly assigned to experimental groups. C57BL/6J mice were obtained from the in-house breeding facility. Details of individual mice involved in the study are listed in Supplementary Table 1.

#### Diets

Mice were typically maintained on a Teklad Global 18% Protein Rodent Diet (Envigo, 2018S). Where indicated in the figure legends, mice were switched to one of the following experimental diets: control amino acid diet (Envigo, TD.220004), serine and glycine deficient amino acid diet (Envigo, TD.220005), based on Handzlik *et al.*^24^. For cohorts in figures 2A and 3A control diet was a modified 5CC7 diet lacking proline, alanine, aspartate, asparagine, and glutamate (TestDiet, 5BQS) or SGD a modified 5CC7 diet lacking proline, alanine, aspartate, asparagine, glutamate, glycine, and serine (TestDiet, 5BQT), based on Maddocks *et al.*^13^. Diet composition is shown in Supplementary Table 2.

#### KL NSCLC model

The mouse model of Kras^LSL-G12D/+^; Lkb1^f/f^ lung cancer^38^ was in C57BL/6J mice (Francis Crick Institute). Lung tumours were induced in male and female mice aged >8 weeks, using 10^7^ plaque-forming units of Adenovirus CMV-Cre (Ad5CMV-Cre) acquired from the University of Iowa Gene Transfer Vector Core (Iowa City, IA). Body weight was typically monitored twice per week since Cre administration or as indicated in figure legends. Mice were humanely euthanized using a pentobarbital injection, upon exhibiting humane endpoint symptoms.

#### KP NSCLC model

The mouse model of Kras^LSL-G12D/+^; Trp53^f/f^ lung cancer was in C57BL/6J mice (Francis Crick Institute). Lung tumours were induced in male and female mice, typically aged >8 weeks, using 10^7^ plaque-forming units of Adenovirus CMV-Cre (Ad5CMV-Cre) acquired from the University of Iowa Gene Transfer Vector Core (Iowa City, IA). Body weight was typically monitored twice per week since Cre administration or as indicated in figure legends. Mice were humanely euthanized using a pentobarbital injection, upon exhibiting humane endpoint symptoms.

#### NSCLC models survival criteria

Mice were considered to have reached the humane endpoint when displaying at least one of the following criteria: body weight loss exceeding 15% of baseline body weight, respiratory distress (including increased or abnormal respiratory rate), or deterioration in body condition, including hunching, piloerection, reduced mobility, or decreased activity.

#### Orthotopic transplantation PDAC mouse model

Orthotopic engraftments of mouse cells were conducted in single caged 8-10-week-old mice. Mice were anesthetized using Isoflurane and a flank incision was made in the left abdominal side. FC1245 (LSL-Kras^G12D/+^;LSL-Trp53^R172H/+^ - KPC) cells (25,000 cells/mouse) were resuspended as a 45 μL suspension of 50% Growth Factor Reduced Matrigel (Corning; 356231) in PBS and injected into the tail region of the pancreas using insulin syringes (29 Gauge). Successful injection was verified by the appearance of a fluid bubble without signs of intraperitoneal leakage. The abdominal wall was sutured with absorbable Vicryl 5-0 suture (Ethicon) using simple continuous pattern and the skin layer was sutured with absorbable Vicryl 6-0 suture (Ethicon) using a half-buried horizontal pattern. The analgesia regime included a subcutaneous injection of meloxicam (5 mg kg^-1^) and buprenorphine (0.1 mg kg^-1^), splash block of bupivacaine (2 mg kg^-1^) locally on muscle layer before closure and administration of meloxicam (5 mg kg^-1^) in apple purée every 24 hours for 3 days post surgery. Mice were also injected with saline (20 μL g^-1^) for fluid support peri-operatively. After 21 days animals were humanely culled using Schedule 1 methods.

#### Subcutaneous PDAC mouse model

2.5x10^5^ cells FC1245 (LSL-Kras^G12D/+^;LSL-Trp53^R172H/+^ - KPC) cells were resuspended in 100 μL PBS and introduced into C57BL/6 mice (males, >9-week old; Francis Crick Institute) by subcutaneous injection on the flank. Tumour measurement and body weight monitoring was performed twice weekly by staff blinded to the experimental outcome. After 14 days animals were humanely culled using Schedule 1 methods. Each mouse was an experimental unit and control mice received a 100 μL PBS injection.

#### Dexamethasone treatment

Dexamethasone 21-phosphate disodium salt (Sigma-Aldrich; D1159) was dissolved in dH2O and administered intraperitoneally (i.p.) at 1mg kg^-1^ daily upon detection of tumours, typically 5 days post tumour cell injection.

#### Metabolic cages

Metabolic phenotyping was performed using the Comprehensive Lab Animal Monitoring System (CLAMS-HC; Columbus Instruments) for open-circuit indirect calorimetry. Mice were single housed for at least 1 week prior to transfer to the metabolic cages. Oxygen consumption (VO₂), carbon dioxide production (VCO₂), food intake and ambulatory activity were continuously monitored using Oxymax software (version 5.71, Columbus Instruments). Data were collected for 5-days beginning immediately upon transfer of the animals to the metabolic cages, at 3 min intervals, under controlled environmental conditions (24°C, 60% relative humidity and 12-h light/dark cycle; lights on from 07:00 to 19:00). Data was exported and analyzed in R Studio (version 2023.06.1+524) using an in-house developed script.

#### Grip Strength

Grip strength was assessed using a grip strength meter (Bio-GS4, Bioseb). Mice were allowed to grasp a metal mesh grid with all four limbs and were gently pulled backward by the base of the tail until they released the grid. Three consecutive measurements were obtained from each mouse with 30 second intervals. The value with the maximum force generated was used for analysis.

#### Plasma collection

Blood was collected with terminal cardiac puncture after S1K culling in Heparin-treated tubes. Following incubation at room temperature for > 20 min, plasma was isolated by centrifugation at 3000 x g for 15 min at 4°C and was stored at −70 °C.

4-hyroxynonenal (4-HNE) plasma measurements were performed using the Universal 4-Hydroxynonenal ELISA Kit (Novus Biologicals; NBP2-66364).

GDF-15 plasma measurements were performed using the Mouse/Rat GDF-15 quantikine ELISA kit (R&D Systems; MGD150).

IL-6 plasma measurements were performed using the Mouse IL-6 quantikine ELISA kit (R&D Systems; M6000B).

Activin-A plasma measurements were performed using the Human/Mouse/Rat Activin A Quantikine ELISA kit (R&D Systems; DAC00B).

Leptin plasma measurements were performed using the Mouse/Rat Leptin Quantikine ELISA kit (R&D Systems; M0B00B).

#### Tissue collection

Liver, tumour, lungs, spleen, visceral and subcutaneous adipose tissue, quadricep and gastrocnemius muscle, samples were typically collected during necropsy dissection, cut into equal parts, which were either snap frozen in liquid nitrogen and stored at -70°C or fixed in 10 % neutral buffered formalin for 24 hours at room temperature before being transferred to 70% ethanol and later paraffin-embedded (FFPE) for immunohistochemistry processing.

### METHOD DETAILS

#### μCT imaging

Lung tumour volume in NSCLC models was measured by micro-CT (μCT) imaging as previously described^62^. Mice were anaesthetised by inhalation of isoflurane and scanned using the Quantum GX2 or GX3 μCT imaging systems (PerkinElmer). X-ray source had 100 kVp and 90 μA, FOV was 25 mm, voxel size 50 μm and the filter (mm) was Cu 0.06 + AI 0.5. Serial lung images were reconstructed and tumour volumes analysed using Analyze (AnalyzeDirect, version 14) as previously described^62^. Whole body scans were performed with μCT using the Quantum GX2 or GX3 μCT imaging systems (PerkinElmer). X-ray source had 100 kVp and 90 μA, FOV was 60 mm, voxel size 120 μm and the filter (mm) was Cu 0.06 + AI 0.5. Whole body serial images were reconstructed and lean, visceral and subcutaneous body compartments were segmented using Analyze (AnalyzeDirect, version 14) software.

#### Tissue protein extraction

Tissue samples were snap-frozen in liquid nitrogen and stored at −70 °C. Samples were homogenized in ice-cold RIPA Lysis and Extraction buffer (ThermoFisher Scientific, 89900) supplemented with protease (cOmplete, EDTA-free Protease Inhibitor Cocktail, Roche, Merck, 4693159001) and phosphatase (PhosSTOP, Roche, Merck, 4906845001) inhibitors. 1 mL of RIPA was used per approximately 50 mg of tissue and samples were homogenised using a Precellys ceramic kit 1.4 mm 2 mL tubes (VWR, 432-3751) and a Precellys 24 homogenizer. Homogenized samples were transferred to Eppendorf tubes, vortexed and incubated on ice for 30 min. Lysates were cleared by centrifugation at 18,000 x g for 10 min at 4°C. Total protein content was quantified by Pierce BCA assay (ThermoFisher Scientific, 23227).

#### Cytokine measurement

Cytokine levels in plasma and liver lysates were measured using the Proteome Profiler Mouse XL Cytokine Array (ARY028, R&D Systems) according to the manufacturer’s instructions. Membranes were imaged using a ChemiDoc MP imaging system (Bio-Rad) and spot intensities were quantified using Fiji software (ImageJ v2.16.0). Cytokine signals were normalized to the positive control reference spots on each membrane and log2FC SGD/control was calculated. Venn diagrams were generated using the online tool Intervene^63^ and pathway enrichment analysis was performed using Metascape^64^.

#### Proteomics

##### Reagents and Chemicals

Ammonium bicarbonate (cat. 40867-50G, Lot No. N0940), phosphoric acid (PCode 102409210), triethylammonium bicarbonate (TEAB) (PCode 102335083), and sodium dodecyl sulfate (SDS) (PCode 1003528300) were purchased from Sigma-Aldrich (St. Louis, MO). Formic acid (FA) 99+% (ref 28905), iodoacetamide (IAA) (cat 122270050), tris(2-carboxyethyl)phosphine hydrochloride (TCEP) (ref PG82080), trypsin protease MS-grade were purchased from Thermo Scientific Inc. (Rockford, IL, USA) (ref 90059). LCMS-grade solvents: water (code W6-4), acetonitrile (ACN) (code A955-212), methanol (code A456-4), were purchased from Fisher Chemical. S-Trap plates were purchased from ProtiFi (Huntington, NY). Evotips were purchased from Evosep (Odense, Danemark).

##### Proteomic sample preparation

Protein digestion in the S-Trap micro column was performed according to the manufacturer’s protocol with some modifications. Briefly, 150 µg protein in lysis buffer (10% SDS in 100 mM TEAB pH 8.5) was reduced by adding TCEP to the protein solution to a final concentration of 5 mM and incubated for 15 minutes at 95 °C. The alkylation was then performed by adding IAA to the reduced samples to a final concentration of 20 mM and incubated for 30 minutes at room temperature in the dark. A final concentration of 1.2% (v/v) phosphoric acid followed by six-fold volume of binding buffer (90%, v/v, methanol in 100 mM TEAB) was next added to the protein solution. After vortexing, the solution was loaded into an S-Trap plate. The solution was removed by spinning the column at 1,000 g for 2 min. The column was washed with 200 μL binding buffer three times. Finally, 125 µL of 0.80 mg/mL trypsin in 50 mM ammonium bicarbonate was added to the S-trap plate and incubated 18 h at 37 °C. Digested peptides were eluted using 80 µL of three buffers consecutively: (1) 50 mM ammonium bicarbonate, (2) 0.2% (v/v) FA in H_2_O, and (3) 50% (v/v) ACN. The elution solutions were collected in a clean 96-well plate and dried under vacuum.

##### Proteomic sample acquisition DIA

Dried samples were resuspended in 0.1% (v/v) FA and 200 ng of tryptic peptides were loaded onto Evotip and analyzed using an Evosep One LC (EVOSEP) connected to a timsTOF Pro (Bruker). The Evosep One method was 60 SPD (21-min gradient, cycle time of 24 min) and the mass spectrometer was operated in DIA-PASEF mode. The DIA-PASEF method consisted of twelve *m/z* windows and a cycle time of 1.37 s. The other mass spectrometer parameters were set as follow: *m/z* range, 262 to 1200, the mobility (1/K0) range was set to 0.60 to 1.60 V.s/cm^2^, and the accumulation and ramp time were 100 ms.

##### Proteomic sample search

The mouse proteins from *Mus musculus* (Uniprot Taxon ID 10090) FASTA file from the Uniprot database was used to generate a spectral library within DIA-NN 1.8 ^65^ (Data-Independent Acquisition by Neural Networks) following a library-free search method. Specific search settings included cysteine carbamidomethylation enabled as a fixed modification, N-terminal methionine excision enabled, maximum missed cleavages 1, min precursor + 2, max precursor + 4, neural network classifier was set to single-pass mode, cross run normalisation was set to retention time dependent and library generation was set to smart profiling. All output was filtered at 0.01 false discovery rate (FDR). Statistical analysis was performed using Perseus software (version 2.0.11) ^66^. Pathway enrichment analysis was performed using Metascape^64^ and volcano plots were generated using the online app VolcaNoseR^67^.

#### Cell culture

Mouse KL (LSL-Kras^G12D/+^;Lkb1^flox^) and KP #01-03 (LSL-Kras^G12D/+^;p53^flox^) NSCLC cell lines were derived from lung tumours of individual C57Bl/6J mice. Mouse FC1245 PDAC cells were derived from tumours on the pancreas driven by *Pdx1-cre*;LSL-Kras^G12D/+^;LSL-Trp53^R172H/+^ in C57Bl/6J mice (originally isolated by Prof. David Tuveson’s lab). Quality control of cell lines was performed by the Francis Crick Institute Cell Science Technology Platform, including mycoplasma testing, STR profiling and species validation. Cells were cultured at 37°C in humidified atmosphere of 5% CO_2_. Cells were cultured in DMEM (ThermoFisher Scientific, Gibco, 41966029) supplemented with 10% FBS (ThermoFisher Scientific, Gibco, A5256701) and penicillin-streptomycin (10,000 units mL^-1^, ThermoFisher Scientific, Gibco, 15140122).

#### Growth curves

For serine and glycine starvation assays, MEM (Thermo Fisher Scientific, Gibco, 21090022) was used as base medium supplemented with L-glutamine (2 mM, Thermo Fisher Scientific, Gibco, 25030024), MEM Vitamin Solution (1%, Merck, Sigma-Aldrich, M6895), 10% dialyzed FBS (Thermo Fisher Scientific, Gibco, 26400044), penicillin–streptomycin (10,000 U ml−1, Thermo Fisher Scientific, Gibco, 15140122) and adjusted with glucose solution (Thermo Fisher Scientific, Gibco, A2494001) to 15.5 mM glucose. This base medium was also supplemented with serine (0.4 mM) and glycine (0.4 mM). 20,000 cells were typically plated on 24-well plates and allowed to adhere overnight. Adherent cells were washed twice with PBS and were fed with assay medium. Medium was replaced with fresh medium every 24 h. Wells were counted (using Casy TT cell counter, Innovatis, Roche Applied Science) at the stated time points, using a ‘time = 0’ plate to calculate relative cell number from time of medium change.

#### Tissue extraction

Tissue samples were snap-frozen in liquid nitrogen and stored at −80 °C. Samples were homogenized in cold (−20 °C) solvent consisting of methanol (Optima LC/MS grade, Fisher Chemical, 10636545) / acetonitrile (Optima LC/MS grade, Fisher Chemical, 10055454) / H_2_O (Optima, LC/MS Grade, Fisher Scientific, 10505904) (50:30:20 v/v). 1 mL of solvent was used per 50 mg of tissue and samples were homogenised using a Precellys ceramic kit 1.4 mm 2 mL tubes (VWR, 432-3751) and a Precellys 24 homogenizer. Homogenized samples were transferred to Eppendorf tubes and 400 μL chloroform was added. Samples were vortexed and transfer to ice-cold water bath sonicator for a 60 min incubation including 3 × 8 minute pulses of sonication. Lysates were then centrifuged at 18,000 x g for 10 min at 4°C. Supernatants were collected to a fresh Eppendorf extraction tube and dried using centrifugal evaporation under vacuum (SpeedVac, RVC 2-33 Cdplus, Martin Christ Gefriertrocknungsanlagen GmbH). Dried pellets were resuspended to a solution of 50 µL chloroform, 150 µL methanol and 150 µL water, vortexed briefly and centrifuged at 18,000 x g for 5 min at 4°C. The upper polar phase was transferred in a new Eppendorf tube, dried and resuspended to 100 μL of acetonitrile (Optima LC/MS grade, Fisher Chemical, 10055454) / H_2_O (Optima, LC/MS Grade, Fisher Scientific, 10505904) (80:20). Samples have been supplemented with 5 μM [^13^C_3,_^15^N]L-Valine (Cambridge Isotope Laboratories, Inc.; CNLM-442-H-0.25) during the extraction process. The apolar lower phase was transferred to a glass vial, dried and resuspended to 300 μL 10 mM ammonium formate (Sigma-Aldrich, 70221) in Butanol (Fisher Scientific; 10223660) : MeOH (Fisher Scientific; 10767665) (1:1 v/v) solution containing Avanti’s EquiSPLASH™ LIPIDOMIX (330731-1EA). Samples were analysed using LCMS.

#### Plasma extraction

Blood was isolated with cardiac puncture after S1K culling in Heparin-treated tubes. Following incubation at room temperature for > 20 min, plasma was isolated by centrifugation at 3000 x g for 15 min at 4°C and was stored at −70 °C. 10 μL of serum was extracted with 30 μL of Methanol (Optima LC/MS grade, Fisher Chemical, 10636545) for 5 min on ice and centrifuged for 10 min at 4 °C and 18,000 x g. Samples were dried using centrifugal evaporation under vacuum (SpeedVac, RVC 2-33 Cdplus, Martin Christ Gefriertrocknungsanlagen GmbH) and partitioned in 50 μL chloroform, 150 μL methanol and 150 μL H_2_O. Samples were centrifuged for 10 min at 4 °C and 18,000 x g. The polar phase was then dried using SpeedVac and was resuspended in 100 μL acetonitrile (Optima LC/MS grade, Fisher Chemical, 10055454) / H_2_O (Optima, LC/MS Grade, Fisher Scientific, 10505904) (80:20 v/v) buffer. Samples have been supplemented with 5 μM [^13^C ^15^N]L-Valine (Cambridge Isotope Laboratories, Inc.; CNLM-442-H-0.25) during the extraction process and analysed using LCMS.

#### Liquid chromatography-mass spectrometry (LCMS)

##### Polar metabolomics analysis

Metabolomics analysis was performed by LCMS using a Orbitrap Exploris 240 mass spectrometer (ThermoFisher) coupled with a Vanquish UHPLC system (ThermoFisher). The chromatographic separation was performed on a Atlantis Premier BEH z-HILIC column (100 × 2.1 mm, 1.7 µm; Waters, Milford, MA, USA). The injection volume was 5 μL, the oven temperature was maintained at 25°C, and the autosampler tray temperature was maintained at 4°C. Chromatographic separation was achieved using a gradient program at a constant flow rate of 300 μL min^-1^ over a total run time of 21 min. Solvent A consisted of 12 mmol L⁻¹ ammonium carbonate (pH 9.0), and solvent B was ACN. The elution gradient was as follows: 95 % solvent B was held for 1 minute, then decreased linearly to 49% over 10 minutes (1–11 min), followed by a 2-minute hold (11–13 min). From 13.01 to 15.00 minutes, solvent B was further decreased to 25 %. At 15.01 minutes, it was returned to 95 % and held until 21 minutes. MS was performed with positive/negative polarity switching with a HESI II probe. MS parameters were as follows: spray voltage 3.5 and 3.0 kV for positive and negative modes, respectively; probe temperature 320°C; sheath and auxiliary gases were 30 and 10 arbitrary units, respectively; and full scan range: 70– 1000 m/z with settings of normalized AGC target (%) and resolution as balanced and high (100 and 120,000), respectively. Mass calibration was performed for both ESI polarities before analysis using the standard ThermoFisher Calmix solution. To enhance calibration stability, lock-mass correction was also applied to each analytical run using ubiquitous low-mass contaminants. Data dependent acquisition (DDA) parameters were the following: resolution 60,000; normalized AGC targeted (%) 100; maximum IT 100 ms; Stepped NCE were: 20, 35, 45. Data were recorded using Xcalibur 4.6 software (ThermoFisher), and the metabolites were identified and quantified by accurate mass and retention time and by comparison to the retention times, mass spectra, and responses of known amounts of authentic standards using TraceFinder 5.1 EFS software (ThermoFisher).

##### Lipidomics analysis

The LC-MS method for lipids was adapted from Amiar et al.^68^ using a Orbitrap Exploris 240 mass spectrometer (ThermoFisher) coupled with a Ultimate 3000 UHPLC system (ThermoFisher). The chromatographic separation was performed on a C18 Zorbax Eclipse Plus column (100 × 2.1 mm, 1.8 µm; Agilent Technologies). Samples were resuspended in 10 mM ammonium formate in Butanol:MeOH (1:1 v/v), and the injection volume was 5 μL, the oven temperature was maintained at 60 °C, and the autosampler tray temperature was maintained at 10°C. Chromatographic separation was achieved at a constant flow rate of 600 μL min^-1^. Solvent A consisted of 10 mmol L⁻¹ ammonium formate, whereas solvent B was H_2_O:acetonitrile:Isopropanol (5:20:75 v/v), containing 10 mmol L⁻¹ ammonium formate. The elution gradient was programmed as follows: solvent B was held at 45% for 2 min, then increased linearly to 100% from 2 to 22 min, followed by a 5 min hold at 100% (22–27 min). At 27.1 min, the initial conditions (45% solvent B) were restored and maintained for 3 min to re-equilibrate the system. The total run time was 30 min. MS was performed with positive/negative polarity switching with a HESI II probe. MS parameters were as follows: spray voltage 3.5 and 2.5 kV for positive and negative modes, respectively; probe temperature 350 °C; sheath and auxiliary gases were 60 and 15 arbitrary units, respectively; and full scan range: 150–2000 m/z with settings of normalized AGC target (%) and resolution as balanced and high (100 and 120,000), respectively. Mass calibration was performed for both ESI polarities before analysis using the standard ThermoFisher Calmix solution. To enhance calibration stability, lock-mass correction was also applied to each analytical run using ubiquitous low-mass contaminants. Data dependent acquisition (DDA) parameters were the following: resolution 60,000; normalized AGC targeted (%) 100; maximum IT 100 ms; Stepped NCE were: 10, 20, 45. Data were recorded using Xcalibur 4.6 software (ThermoFisher). The .raw files were imported into Progenesis QI v3.1.9211.37750 (Nonlinear Dynamics, Waters™) which was used for automated retention time alignment and peak detection. Peak picking utilized the software’s automatic sensitivity parameter method using the default (level 3) noise estimation algorithm for ion detection. Sample data were normalized against the total ion abundance. Additionally, features were also annotated based on accurate mass match with the LipidMatch R script v 4.2^69^.

For metabolomics and lipidomics heatmaps and volcano plots were generated using Metaboanalyst 6.0^70^. PCA plots were generated using SIMCA (Sartorius, version 17).

#### Histology and Immunohistochemistry

Tissues were fixed in 10% neutral buffered formalin for 24 hours. The fixed tissues were processed using Sakura Tissue-Tek VIP® 6 AI Tissue Processor and paraffin embedded. 3μm sections were cut on glass slides. The slides were baked at 60°C for 1 hour and then deparaffinized in xylene and rehydrated using a series of graded industrial methylated spirits solutions and distilled water. For H&E staining the slides were stained using Tissue-Tek Prisma® Plus Automated Slide Stainer. For immunohistochemistry, heat-mediated antigen retrieval was performed using pH 6 citrate buffer for 20 minutes in a microwave. Endogenous peroxidase blocking was performed by incubating the slides in Bloxall (SP-6000-100, Vector) for 10 minutes at room temperature. The slides were incubated with 2.5% Normal Horse Serum for 30 minutes at room temperature. Primary antibodies Anti-Ki67 (1:500, Abcam ab15580, LOT 1063774-1), Anti-F4/80 (1:2500, Abcam ab300421, LOT 1043287-14) and Ly-6G (1:100, Cell Signalling 87048, LOT 6) were applied for 1 hour at room temperature. The slides were incubated in ImmPRESS anti-rabbit Polymer Reagent (Vector MP-7401) for 30 minutes at room temperature. Slides were incubated with 3,3-diaminobenzidine (DAB) chromogen (SK-4105, Vector) for 10 minutes at room temperature. The slides were then counterstained using Harris Haematoxilin, dehydrated, cleared and mounted using Sakura Tissue-Tek Prisma automated stainer. The slides were scanned using Zeiss Axioscan 7 Slide Scanner using a 10X brightfield scanning profile. QuPath (version 0.5.1)^71^ was used for IHC analysis.

#### Statistics and reproducibility

All datasets were analysed and plotted using Prism 11 (v11.0.0; GraphPad Software) unless otherwise stated. Differences between groups were tested for normal distribution and analysed using the appropriate statistical test, as mentioned in each figure legend. Error bars represent s.d., unless otherwise stated.

### Data availability

Associated raw data are provided as supplementary source files associated with each main or supplementary figure. Original datasets, analyses and methodological details are available upon request. Information regarding reagents can also be found in supplementary table 3.

## SUPPLEMENTARY FIGURES & LEGENDS

**Supplementary Figure 1.**
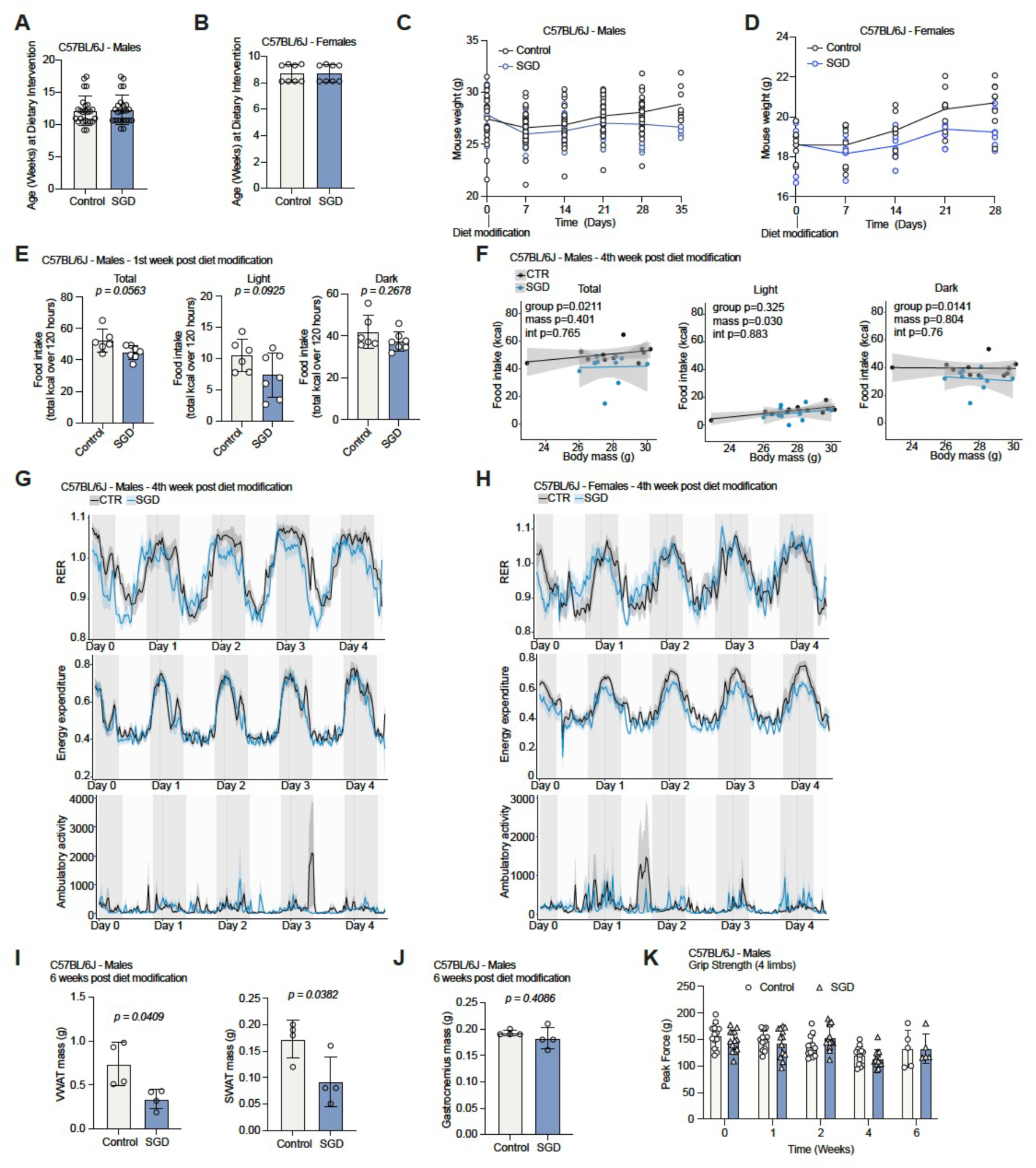
SGD diet decreases body weight and adiposity while preserving skeletal mass in WT C57Bl/6J mice. (A) Age (weeks old) of C57Bl/6J WT male mice at the time of enrolment. Data are from n = 25 Control and 24 SGD mice. (B) Age (weeks old) of C57Bl/6J WT female mice at the time of enrolment. Data are from n = 8 Control and 8 SGD mice. (C) Weight trajectories of C57Bl/6 WT male mice from diet enrolment till collection. Data are from n = 25 Control and 24 SGD mice. (D) Weight trajectories of C57Bl/6 WT female mice from diet enrolment till collection. Data are from n = 8 Control and 8 SGD mice. (E) Food intake (total kcal over 120 hours) for the total period, light phase and dark phase (as indicated) of C57Bl/6J WT male mice in control or SGD diet during the 1^st^ week post diet modification. Values are mean±s.d. from n = 6 control and n = 7 SGD mice. (F) Food intake (kcal) over body mass (g) of C57Bl/6 WT male mice in control (CTR) or SGD diet 4 weeks post diet modification. Values are from n = 11 Control and 11 SGD mice for the total period, light phase and dark phase (as indicated). Solid lines represent fitted values from ANCOVA analysis with group as a fixed factor and body mass as a covariate; shaded regions denote 95% confidence intervals. (G) Timeseries of RER (top), Energy expenditure (middle) and Ambulatory activity (bottom) of C57Bl/6J WT male mice in control (CTR) or SGD diet 4 weeks post diet modification for the total duration of the recording. Values are average from n = 11 Control and 11 SGD mice with line shading indicating ±s.e.m. (H) Timeseries of RER (top), Energy expenditure (middle) and Ambulatory activity (bottom) of C57Bl/6J WT male mice in control (CTR) or SGD diet 4 weeks post diet modification for the total duration of the recording. Values are average from n = 11 Control and 11 SGD mice with line shading indicating ±s.e.m.. (I) VWAT and SWAT mass (gr) of C57Bl/6J WT male mice in control or SGD diet 6 weeks post diet modification. Values are mean±s.d. from n = 4 Control and 4 SGD mice. (J) Gastrocnemius mass (gr) of C57Bl/6J WT male mice in control or SGD diet 6 weeks post diet modification. Values are mean±s.d. from n = 4 Control and 4 SGD mice. (K) Grip strength force (g) of C57Bl/6 WT male mice in control (CTR) or Ser/Gly-deficient diet (SGD) over time post diet change. Values are mean±s.d. from n = 13 Control and 13 SGD mice (weeks 0-4) and 5 Control, 5 SGD mice at week 6. Statistical significance was assessed by two-tailed Welch’s t-test (I, J) and two-tailed unpaired t-test with Benjamini-Hochberg correction for multiple comparisons (E).

**Supplementary Figure 2.**
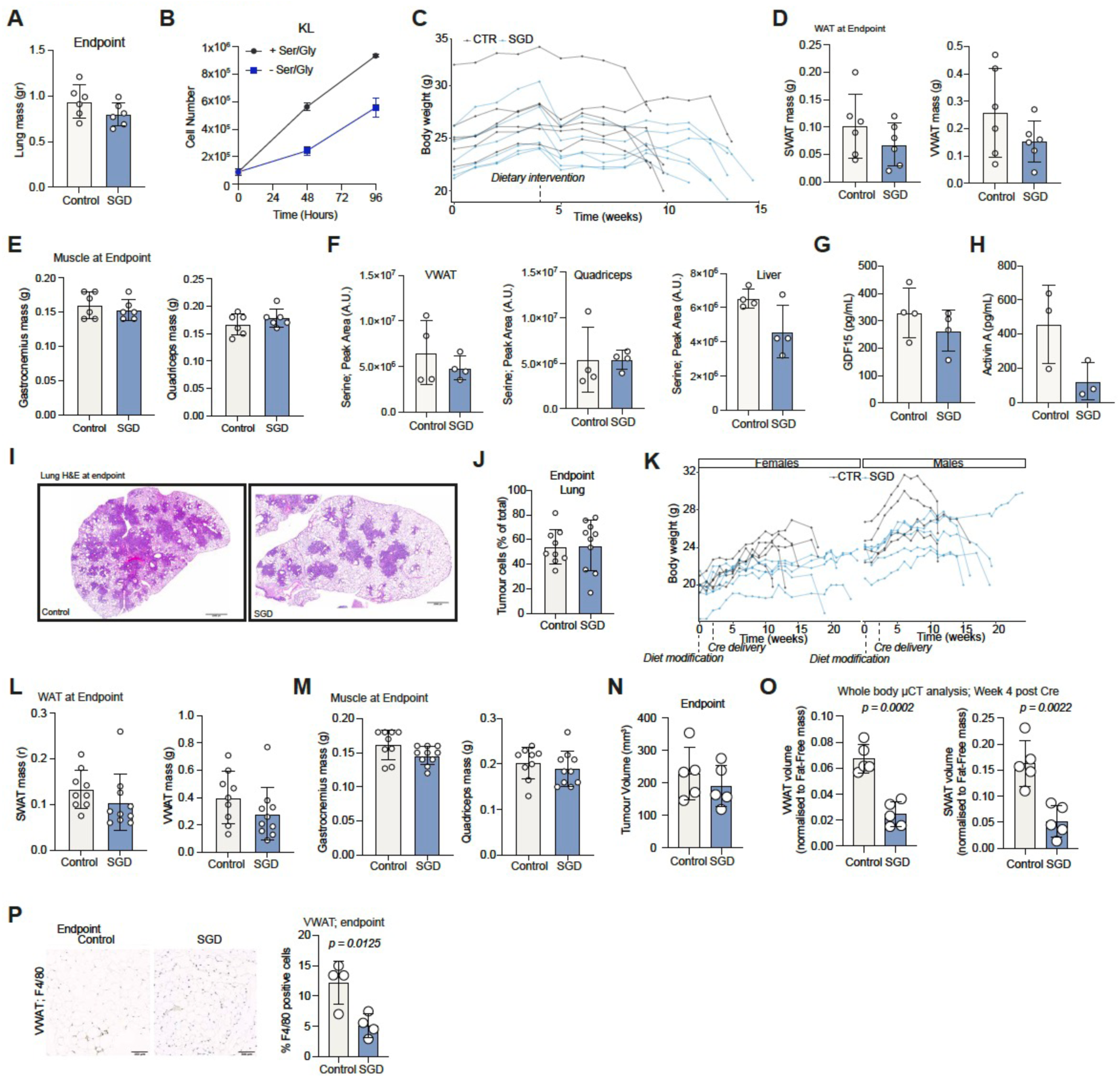
SGD diet suppresses tumour progression and delays cachexia in Kras^G12D/+^;Lkb1^f/f^ NSCLC. (A) Lung mass (gr) of KL male mice treated with control or SGD diet at endpoint. Values are mean±s.d. from n = 6 control and 6 SGD mice. (B) Proliferation curves of KL cells ±Serine/Glycine. Values are mean±s.d. of triplicate wells. (C) Weight trajectories of KL mice treated with control (n = 6) or SGD diet (n = 6). (D) SWAT (left) and VWAT (right) mass of mice from (B) at endpoint. Values are mean±s.d. from n = 6 control and 6 SGD mice. (E) Gastrocnemius (left) and Quadriceps (right) mass of mice from (B) at endpoint. Values are mean±s.d. from n = 6 control and 6 SGD mice. (F) Serine levels in VWAT, Quadriceps and liver lysates of mice from (B) as indicated. Values are mean±s.d. from n = 4 control and 4 SGD KL mice. (G) Plasma GDF15 levels in mice from (B) at endpoint. Values are mean±s.d. of n = 4 control and 4 SGD mice. (H) Plasma Activin A levels in mice from (B) at endpoint. Values are mean±s.d. of n = 3 control and 3 SGD mice. (I) Representative pictures of H&E-stained lungs at endpoint from KL mice preconditioned with control or SGD diet. Scale bars 1000 μm. (J) Quantification of tumour burden in mice from (I). Values are mean±s.d. of n = 9 control (5 female & 4 male) and n = 10 SGD mice (5 female and 5 male). (K) Weight trajectories of KL mice preconditioned with control (n = 9) or SGD diet (n = 10). (L) SWAT and VWAT mass of mice from (K) at endpoint. Values are mean±s.d. from n = 9 control and 10 SGD mice. (M) Gastrocnemius and Quadriceps mass of mice from (K) at endpoint. Values are mean±s.d. from n = 9 control and 10 SGD mice. (N) Tumour volume from μCT quantification prior to endpoint for each mouse. Data are from (Figure 2N) shown as bar plots and selecting only mice with similar tumour burden at endpoint. (O) VWAT (left) and SWAT (right) depots (μCT quantification) at week 4 post Cre administration, for mice with similar tumour burden at endpoint. Values are mean±s.d. from n = 5 control and 5 SGD mice. Statistical significance was assessed by two-tailed unpaired Welch’s t-test. (P) Representative pictures (left) and quantification (right) of F4/80-stained VWAT of mice from (B) at endpoint. Scale bars, 100 μm. Values are mean±s.d. from n = 4 control and n = 4 SGD mice. Statistical significance was assessed by two-tailed unpaired t-test.

**Supplementary Figure 3.**
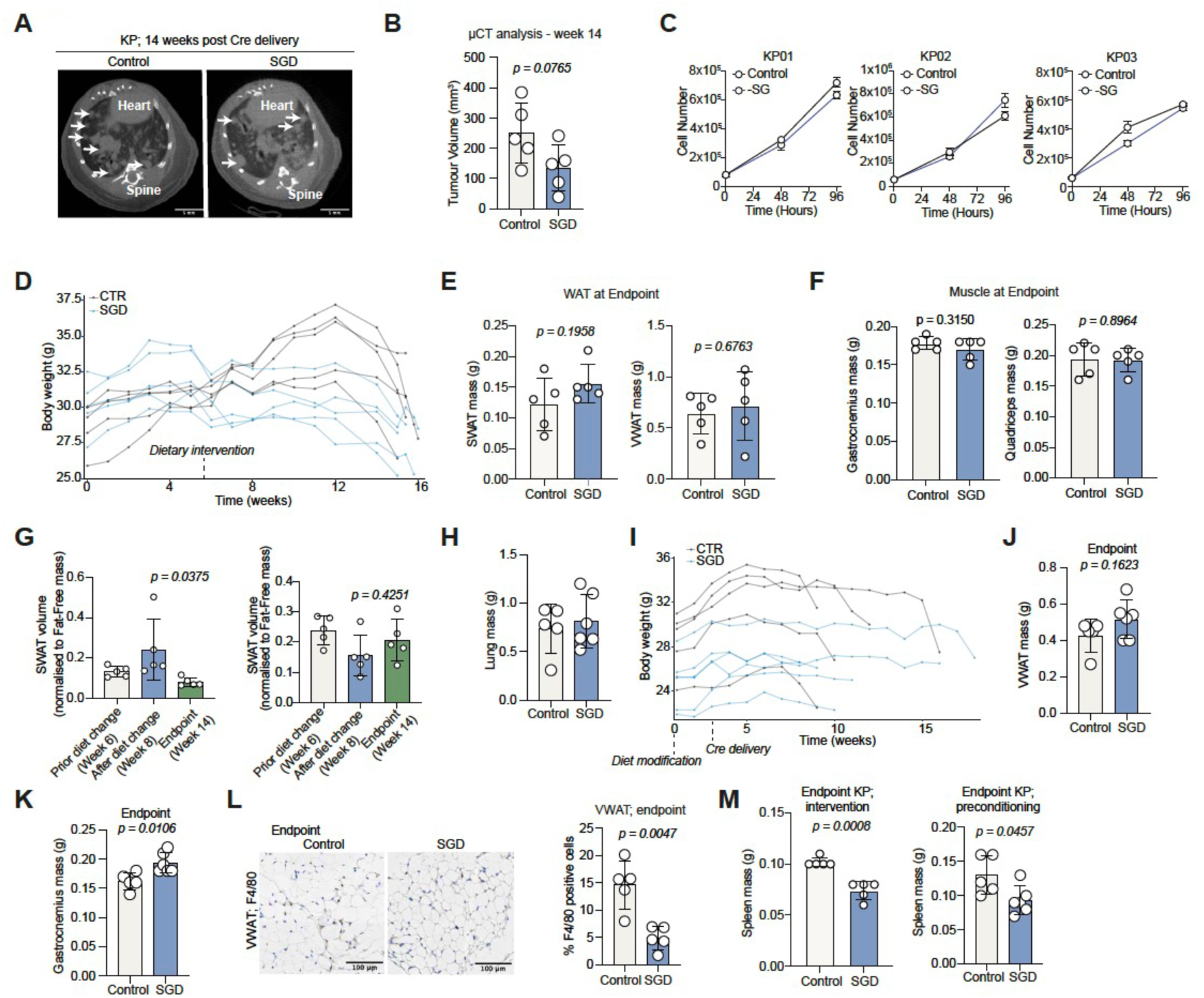
SGD diet delays cachexia independently of tumour growth in Kras^G12D/+^;p53^f/f^ mutant NSCLC. (A) Representative μCT image showing lung tumours in KP mice 14 weeks post Cre delivery treated with control or SGD diet. Scale bars, 5mm. (B) Tumour volume (μCT quantification) at week 14 post Cre delivery. Data are from (B) shown as a bar plot with mean±s.d. values from n = 5 control and n = 5 SGD mice. (C) Proliferation curves of KP01, KP02, KP03 cells in complete (control) or SGD cell culture medium. Values are mean±s.d. of triplicate wells. (D) Weight trajectories of KP mice from (B). (E) SWAT (left) and VWAT (right) mass of mice from (B) at endpoint. Values are mean±s.d. of n = 5 control and 5 SGD mice. (F) Gastrocnemius (left) and Quadriceps (right) mass of mice from (B) at endpoint. Values are mean±s.d. of n = 5 control and 5 SGD mice. (G) SWAT volume (ratio to Fat-Free mass) of control (n = 5) (left panel) and SGD (n = 5) (right panel) KP mice before (week 6), after (week 8) diet modification and before endpoint. (H) Lung mass of of KP mice preconditioned with control or SGD diet at endpoint. Values are mean±s.d. from n = 5 control and n = 6 SGD mice. (I) Weight trajectories of KP mice from (I). (J) VWAT mass of mice from (I) at endpoint. Values are mean±s.d. of n = 5 control and 6 SGD mice. (K) Gastrocnemius mass of mice from (I) at endpoint. Values are mean±s.d. of n = 5 control and 6 SGD mice. (L) Representative pictures (left) and quantification (right) of F4/80-stained VWAT of mice from (B) at endpoint. Scale bars, 100 μm. Values are mean±s.d. from n = 5 control and n = 5 SGD mice. Statistical significance was assessed by two-tailed unpaired t-test. (M) Spleen mass of KP mice at endpoint of mice from (B) (left) and (I) (right) . Values are mean±s.d. of n = 5 control and 5 SGD mice (left) and n = 5 control and n = 6 SGD mice (right). Statistical significance was assessed by two-tailed Welch’s t-test (B,E,F,J,K,L,M) and ordinary one-way ANOVA with Tukey’s multiple comparisons test (H).

**Supplementary Figure 4.**
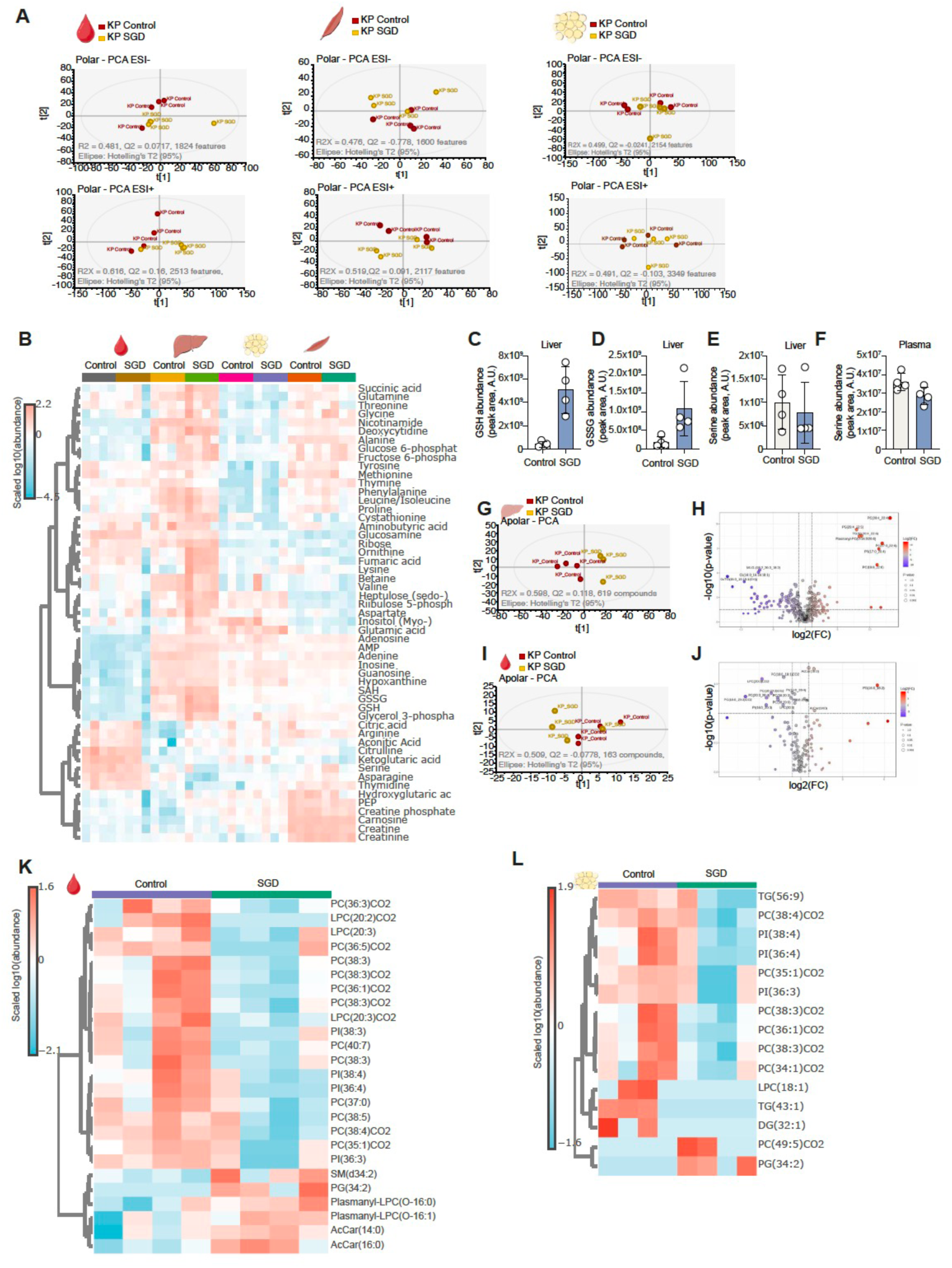
SGD diet rewires hepatic metabolism and reduces systemic oxidative stress in KP NSCLC. (A) PCA plots of LCMS analysis of plasma (left), gastrocnemius (middle) and VWAT (right) polar extracts from control or SGD-treated KP mice. (B) Heatmap, abundance of identified polar metabolites in plasma, liver, VWAT and gastrocnemius lysates from control or SGD treated KP mice. (C) GSH levels in liver lysates of KP mice treated with control or SGD diet. Values are mean±s.d. from n = 4 control and n = 4 SGD mice. (D) GSSG levels in liver lysates of KP mice treated with control or SGD diet. Values are mean±s.d. from n = 4 control and n = 4 SGD mice. (E) Serine levels in liver lysates of KP mice treated with control or SGD diet. Values are mean±s.d. from n = 4 control and n = 4 SGD mice. (F) Serine levels in plasma of KP mice treated with control or SGD diet. Values are mean±s.d. from n = 4 control and n = 4 SGD mice. (G) PCA plots of LCMS analysis of liver apolar extracts from control or SGD-treated KP mice. (H) Volcano plot of identified lipid species in liver lysates from control or SGD-treated KP mice. (I) PCA plots of LCMS analysis of plasma apolar extracts from control or SGD-treated KP mice. (J) Volcano plot of identified lipid species in plasma lysates from control or SGD-treated KP mice. (K) Heatmap, abundance of identified lipid species in apolar plasma extracts from control or SGD treated KP mice. Only top 25 statistically significant identified lipid species are displayed. (L) Heatmap, abundance of identified lipid species in apolar VWAT extracts from control or SGD treated KP mice. Only top 15 statistically significant identified lipid species are displayed.

**Supplementary Figure 5.**
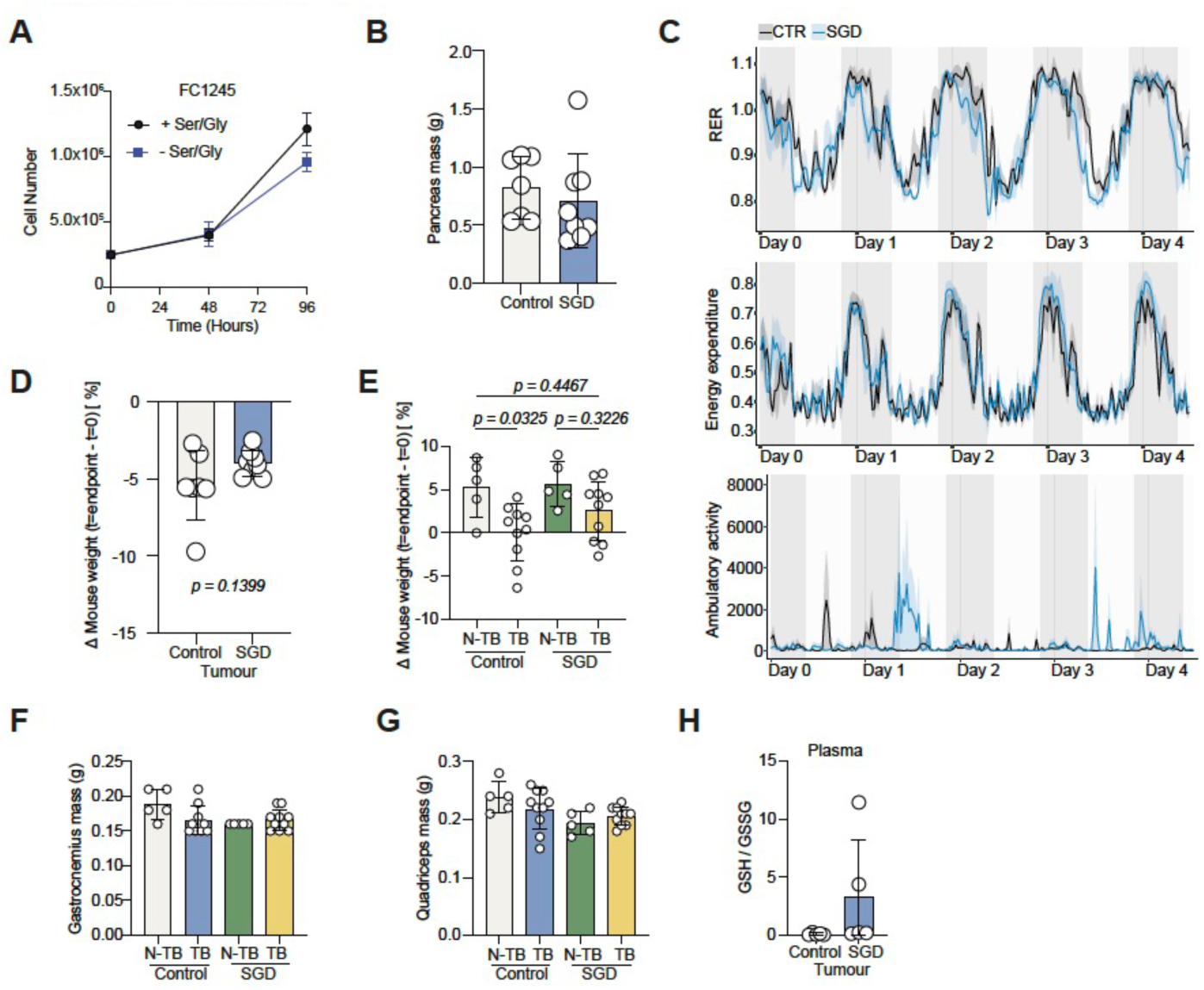
SGD diet improves host responses in SGD-resistant models of PDAC. (A) Proliferation curves of FC1245 KPC cells in complete (control) or Serine/Glycine-deficient cell culture medium. Values are mean±s.d. of triplicate wells. (B) Pancreas mass (gr) of orthotopic KPC PDAC mice preconditioned with control or SGD diet. Values are mean±s.d. from n = 7 tumour control diet and n = 8 tumour SGD diet. (C) Timeseries of RER (top), Energy expenditure (middle) and Ambulatory activity (bottom) of orthotopic KPC PDAC mice preconditioned with control or SGD diet. Values are average from n = 4 Control and 4 SGD mice with line shading indicating ±s.e.m. (D) Weight difference (%) of endpoint (minus tumour weight) vs starting weight (prior tumour implantation) of mice from (B). Values are mean±s.d. from n = 7 control and n = 8 SGD diet mice. (E) Weight difference (%) of endpoint (minus tumour weight) vs starting weight (prior tumour implantation) of KPC subcutaneously transplanted mice preconditioned with control or SGD diet. Values are mean±s.d. from n = 5 control diet WT, n = 10 control diet KPC transplanted, n = 5 SGD diet WT and n = 10 SGD diet KPC transplanted mice. (F) Gastrocnemius mass of mice from (E). Values are mean±s.d. from n = 5 control diet WT, n = 10 control diet KPC transplanted, n = 5 SGD diet WT and n = 10 SGD diet KPC transplanted mice. (G) Quadriceps mass of mice from (E). Values are mean±s.d. from n = 5 control diet WT, n = 10 control diet KPC transplanted, n = 5 SGD diet WT and n = 10 SGD diet KPC transplanted mice. (H) GSH/GSSG ratio in plasma of mice from (E). Values are mean±s.d. from n = 5 control and n = 5 SGD mice. Statistical significance was assessed by two tailed unpaired Welch’s t-test (D) and ordinary one-way ANOVA with Tukey’s multiple comparisons test (E).

**Supplementary Figure 6.**
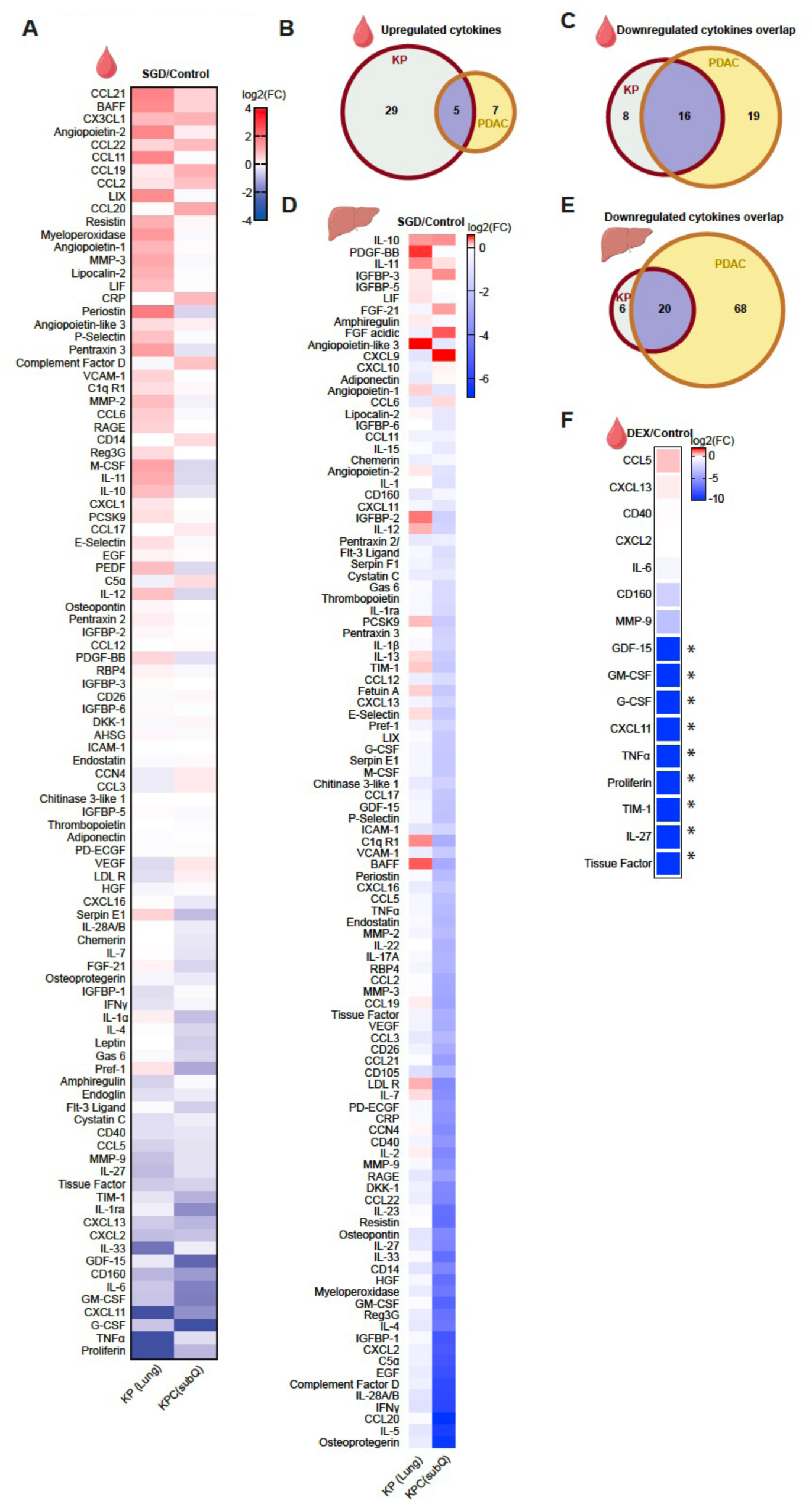
SGD diet suppresses acute systemic inflammatory signalling associated with cachexia. (A) Heatmap, plasma cytokine analysis in SGD-treated KP and PDAC subcutaneous models. (B) Venn diagram, overlap of upregulated cytokines in plasma of SGD-treated KP and PDAC subcutaneous models. (C) Venn diagram, overlap of downregulated cytokines in plasma of SGD-treated KP and PDAC subcutaneous models. (D) Heatmap, liver cytokine analysis in SGD-treated KP and PDAC subcutaneous models. (E) Venn diagram, overlap of downregulated liver in plasma of SGD-treated KP and PDAC subcutaneous models. (F) Heatmap, plasma cytokine analysis in DEX-treated mice with PDAC subcutaneous tumours. (*, undetectable in DEX lysates).

